# Effects of calibrated blue–yellow (–S+[L+M], +S–[L+M]) changes in light on the human circadian clock

**DOI:** 10.1101/2023.08.01.551458

**Authors:** Christine Blume, Christian Cajochen, Isabel Schöllhorn, Helen C. Slawik, Manuel Spitschan

## Abstract

Evening exposure to short-wavelength light can acutely affect the circadian clock located in the suprachiasmatic nuclei, sleep, and alertness. The intrinsically photosensitive retinal ganglion cells (ipRGCs) expressing the photopigment melanopsin are thought to be the primary drivers of these effects. Much less is known about the contribution of the colour-sensitive cones. Using calibrated silent-substitution changes in light colour along the blue-yellow axis, we investigated whether mechanisms of colour vision affect the human circadian system and sleep. In a 32.5-h repeated within-subjects protocol, 16 healthy participants (8 women, 18-35 years old) were exposed to three different light scenarios for 1 h starting 30 min after habitual bedtime: a control condition (“background”, 93.5 photopic lux), intermittently flickering yellow-bright light (1 Hz, 30s on-off, 123.5 photopic lux), and intermittently flickering blue-dim light (1 Hz, 30s on-off; 67.0 photopic lux). Importantly, there was no difference in melanopsin excitation (163.2±2.1 lux melanopic EDI) between the three lighting conditions, allowing us to determine the unique contribution of the blue-yellow colour system. Our analyses did not yield conclusive evidence for differences between the three lighting conditions regarding circadian melatonin phase delays, melatonin suppression, subjective sleepiness, psychomotor vigilance, or sleep. Thus, in this study, we found no evidence that evening light changing along the blue-yellow dimension under moderate light levels typical for room illumination has a major impact on the human circadian clock or sleep. Our work underscores the previously demonstrated primary role of melanopsin-containing ipRGCs in mediating these effects.

## Introduction

### Effects of light on circadian physiology

Light profoundly affects human physiology and behaviour via the retinohypothalamic pathway that relays information from the retina to the circadian pacemaker in the suprachiasmatic nuclei^1^. In humans, this pathway has been functionally mapped out by examining acute responses to evening or night-time light exposure on melatonin secretion, or its circadian phase-shifting effects. Both neuroendocrine (i.e., evening and nocturnal melatonin suppression^2–4^) and circadian-phase shifting responses^5, 6^ to light are primarily driven by the photopigment melanopsin expressed in subset of so-called intrinsically photosensitive retinal ganglion cells (ipRGCs)^7^.

### Melanopsin, cone and rod inputs to circadian phototransduction

However, melanopsin-expressing ipRGCs also receive input from the classical retinal photoreceptors, the cones (active during daylight light levels) and rods (sensitive to dim light but saturated in daylight). Primate ipRGCs receive excitatory synaptic input from the L and M cones and the rods, and inhibitory inputs from the S-cones^8^ mediated by an S-cone amacrine cell^9^ (Figure **1*a***). In humans, evidence for this inhibitory input to ipRGCs has been found in the pupillary light response, where S-cone input inhibits the melanopsin-induced pupil constriction^10, 11^. A recent direct test for an S-cone involvement in melatonin suppression in the evening has led to the conclusion that they do not contribute to acute neuroendocrine responses in constant light^12^ at ∼170 lux, and there is evidence that cones are not necessary for circadian and neuroendocrine responses to light as some visually blind individuals suppress melatonin in response to light^13, 14^. Thus, under continuous lighting conditions, circadian and neuroendocrine responses to light are largely driven by melanopsin^4, 6^.

**Figure 1.**
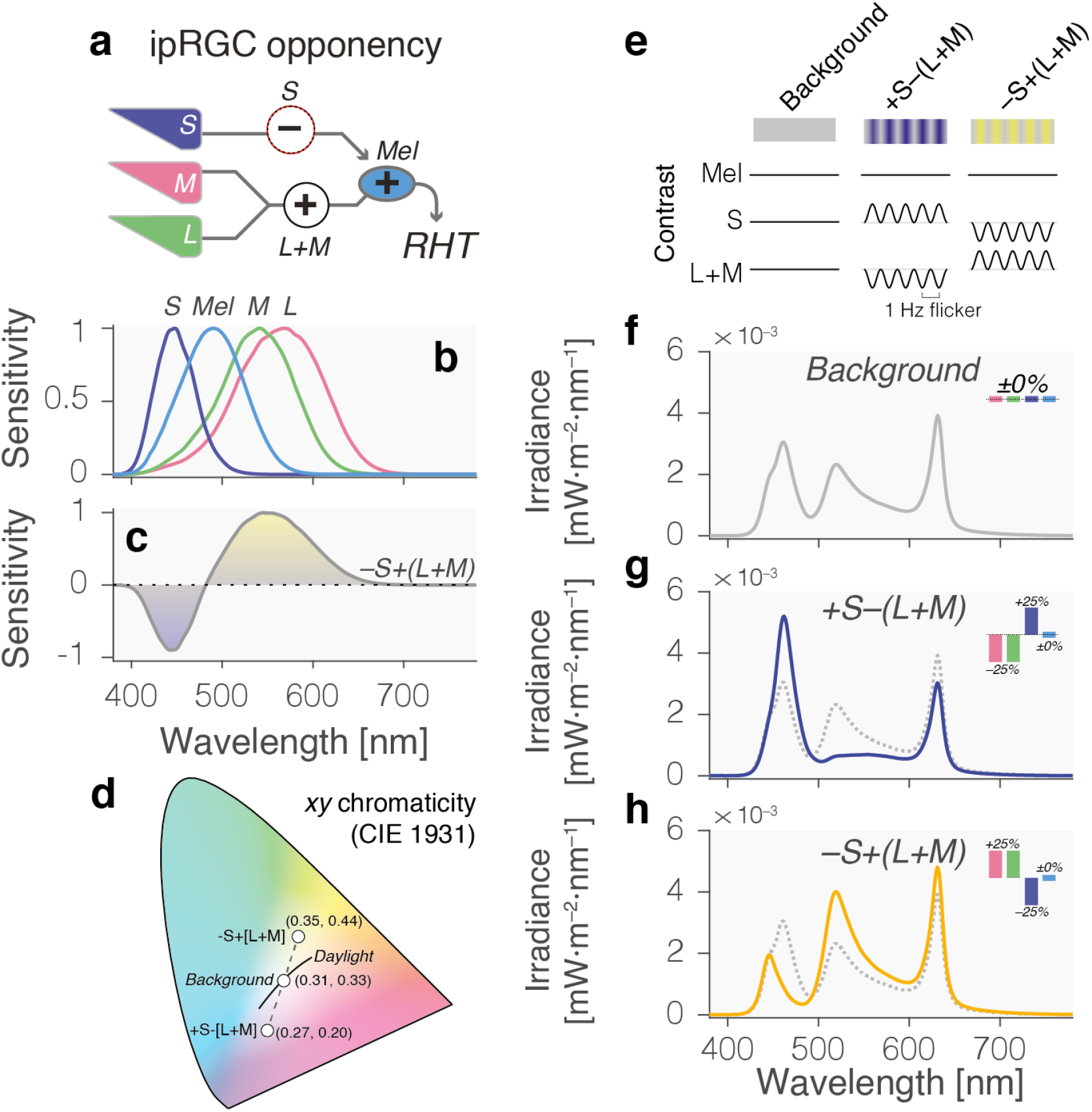
S opponent influences into ipRGCs and selective stimuli to target these pathways. ***a*** The cone inputs to the melanopsin-containing intrinsically photosensitive retinal ganglion cells (ipRGCs) in the human retina combine light information in an opponent, subtractive fashion, pitting signals from the S-cones against signals a joint L and M cone signal encoding luminance (schematic diagram, simplifying underlying the underlying anatomy). ***b*** The spectral sensitivities of the L, M and S-cones have distinct peaks (λ_max_) but are overlapping. ***c*** The joint spectral sensitivity of the opponent –S+(L+M) channel, showing distinct wavelength regions yielding positive vs. negative activations. ***d*** Chromaticity diagram (CIE 1931 *xy* chromaticity) showing the chromaticity coordinates of the background condition (corresponds to colour temperature of 6,500K daylight [D65]), the two modulation spectra and the daylight locus (daylight spectra between 4,000K and 25,000K). ***e*** Overview of stimulus conditions and its contrast properties Stimuli are unipolar sinusoidally flickering excursions from the background in the direction of the +S–(L+M) and –S+(L+M) poles of the blue-yellow channel. ***f*** Spectral irradiance distribution of the constant background, keeping excitation constant for L, M and S-cones and melanopsin. *Inset*: Contrast for the L, M, and S-cones, and melanopsin for the modulation spectrum against the background spectrum. ***g*** Spectral irradiance distribution of the +S–(L+M) stimulus, biasing S-cones over luminance (blue solid line) against the background (dashed grey line). *Inset*: Contrast for the L, M, and S-cones, and melanopsin for the modulation spectrum against the background spectrum. ***h*** Spectral irradiance distribution of the –S+(L+M) stimulus, biasing S-cones over luminance (blue solid line) against the background (dashed grey line). *Inset*: Contrast for the L, M, and S-cones, and melanopsin for the modulation spectrum against the background spectrum.

### Prior evidence for S-cone opponent circuitry

In parallel to the physiological opponency of cone signals established using electrophysiological recordings in the primate retina^15^, there is ample evidence that human image-forming colour vision is also organised according to colour-opponent channels. Signals from the three cone classes with overlapping but distinct spectral sensitivities (Figure **1*b***) are recombined into three channels^16, 17^: an additive combination of L and M cones (L+M; luminance), an opponent, subtractive combination of L and M cones (L–M; red-green), and an opponent combination pitting S-cones against luminance (S–[L+M]; blue-yellow; Figure **1*c***). These post-receptoral channels form the basic dimensions of human colour vision, also called the cardinal directions of colour space^18^. It has previously been suggested, but not directly tested that this circuit may participate in the suppression of nocturnal melatonin secretion by light^19^.

### Dawn and dusk changes encoded by S-cone opponent circuitry

As the transition between day and night, twilight represents a key change in the light environment with changes in light intensity being largest during dusk and dawn^20^. Importantly, however, the spectral composition of environmental illumination also changes, with a boost in short-wavelength illumination relative to long-wavelength light^20, 21^. This signal has long been hypothesised to be informative for the timing of activity and circadian photoentrainment, although this precise relationship has not yet been established^21–23^. In general, a colour-opponent system pitting short-wavelength signals against long-wavelength signals is well-suited to pick up the spectral changes at dawn and dusk^22^, but whether there is a dedicated colour-opponent input into the human circadian clock has presently not been demonstrated.

### Evidence for an +S–L opponent channel in murine circadian phototransduction

Recently, it was reported that in mice, stimuli defined along the post-receptoral axis pitting murine S-cones against murine L-cones differ in their circadian effect, with ‘yellow’ and bright light (with high L-cone activation and low S-cone activation, activating a +S–L opponent channel) induced a stronger circadian phase-shift, as assessed by behavioural responses, than ‘blue’ and dim light (with lower L-cone activation and high S-cone activation, activating an –S+L opponent channel)^24^. This is somewhat counter-intuitive in the light of prior research as many studies in humans have shown that short-wavelength (enriched) light exerts stronger effects on the circadian system^1, 19, 25, 26^. Although it is unknown whether this effect translates to humans, primate ipRGCs are characterised by an +S–(L+M) opponent organisation as well^8, 9^, thereby providing a homologous substrate for the effects seen in mice. Based on these converging lines of evidence (see Table **3** for a summary of the extant literature), we therefore hypothesized that a similar effect can be found in the circadian system of humans.

In this *Registered Report*, we examined the circadian phase-shifting effects of evening light exposure incorporating calibrated changes along the +S–[L+M], blue-yellow dimension of human vision and its effects on sleep in the ensuing night. Specifically, we examined circadian phase shifts in the melatonin rhythm in response to 1-hour light exposure to three different scenarios in a 32.5-hour, within-subjects in-laboratory protocol under controlled lighting and temperature conditions as well as its effects on acute melatonin suppression, subjective and objective sleepiness, visual comfort, psychomotor vigilance, sleep onset and slow-wave activity during sleep (cf. Figure **2**).

**Figure 2.**
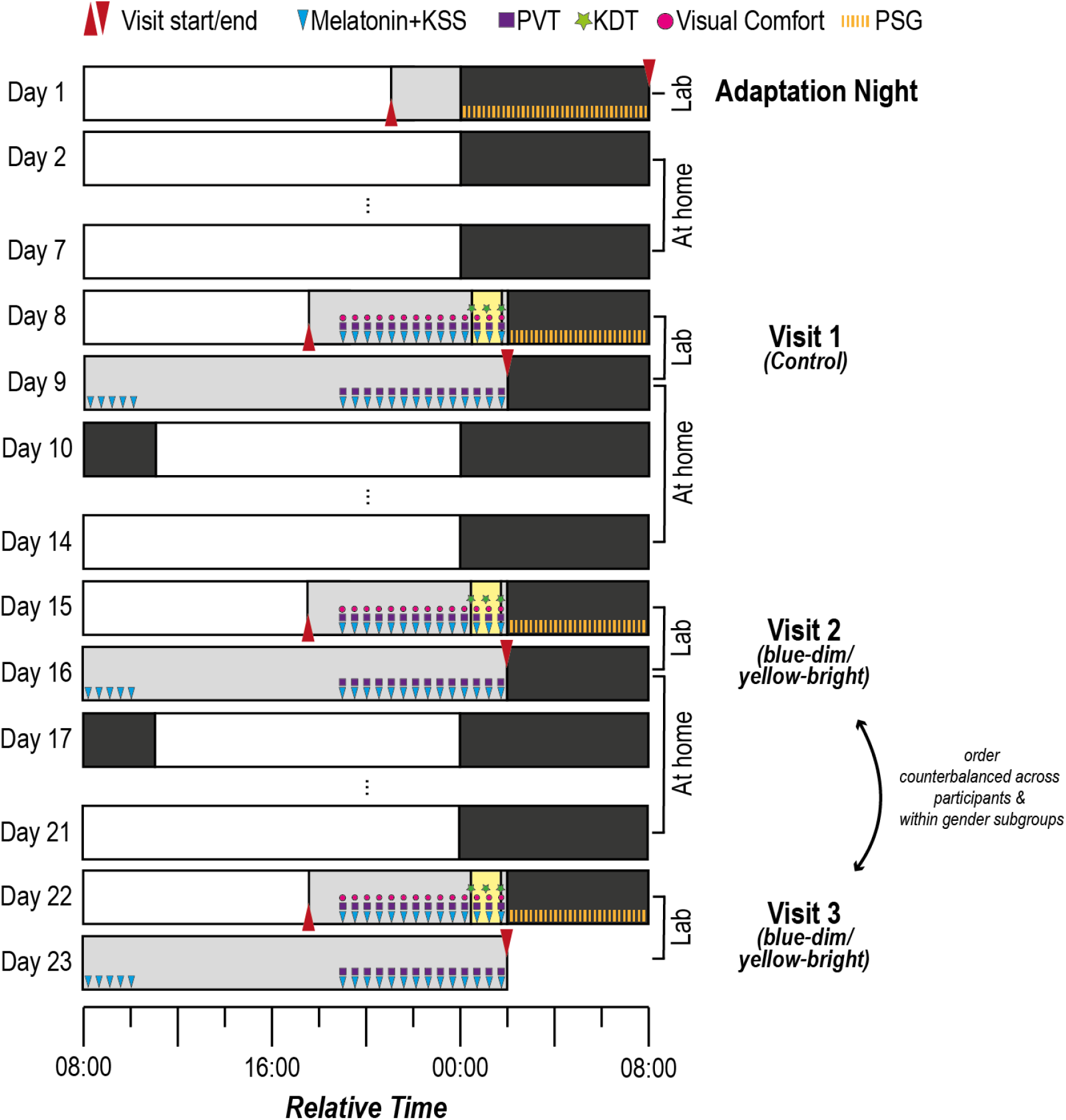
Experimental protocol for within-subjects assessment of light-exposure conditions. Participants engage in a circadian stabilisation protocol for a total of 23 days, during which they visit the laboratory for a total of four times: 1x adaptation night (Day 1) and three **32.5-h**our protocol visits to assess the effect of the light exposure (1 h each starting 30 min after habitual bedtime, as indicated by the yellow boxes) conditions on the human circadian clock. The first 32.5-hour visit (Protocol 1 on Days 8 and 9) was always the constant background (“control”) light, while the subsequent two visits were balanced between participants (*n*=16 in total, 8 women and 8 men). For the experimental visits, volunteers arrived at the lab 6.5 h prior to their habitual bedtime. Melatonin sampling and assessments of subjective sleepiness (with the Karolinska Sleepiness Scale, KSS), psychomotor vigilance (with the psychomotor vigilance task, PVT) as well as assessments of visual comfort in 30 min intervals started 5 hours prior to habitual bedtime. Before, during, and after light exposure, we also assessed objective sleepiness with the Karolinska Drowsiness Test (KDT). Subsequently, participants went to bed for a 6-h sleep opportunity. Following wake-up, another five melatonin samples as well as KSS assessments were obtained. During the second evening, PVT and KSS measurements were performed as on evening one. During all nights, polysomnography (PSG) was recorded. The figure shows the relative time for a participant whose habitual bed time is at midnight. Gray boxes indicate dim light (i.e., mostly < 8 lux).

Stimuli were presented in wide-field, Newtonian (no artificial pupil) viewing without pharmacological dilation as sinusoidally flickering modulations (1 Hz, 30 sec on, 30 sec off to avoid adaptation) either favouring S-cones over luminance (+S–[L+M]; blue-dim) or luminance over S-cones (–S+[L+M]; yellow-bright) with no change in melanopsin activation around a background mimicking daylight (D65, 6500K; planned 100 lux [measured: 93.5 lux]). Additionally, we also examined the response of light to this constant background.

### Primary, confirmatory hypothesis

The primary outcome variable of this study was the circadian phase shift in salivary melatonin onset in the three conditions, constant light (always condition 1), blue-dim flickering, and yellow-bright flickering, which were administered to each participant in a within-subjects design (see Tables **1 and 2** for a description of the outcome variables). **We specifically hypothesized that yellow-bright flickering would induce a larger circadian phase shift than blue-dim changes and that flickering generally induces larger shifts than constant light** (yellow-bright > blue-dim > constant background; confirmatory test C_1_; see Table **1**).

**Table 1.** Design Table.

| Question | Hypothesis | Sampling plan | Analysis Plan | Interpretation given to different outcomes |
| --- | --- | --- | --- | --- |
| **Primary Outcome** |  |  |  |  |
| **Dim-light melatonin onset (DLMO) (C1)** | The difference in DLMO (in minutes) between evenings 1 and 2 of the protocol are larger in the yellow-bright condition than in the blue-dim condition. The DLMO difference in both conditions is larger than in the constant light condition.<br>Effect of interest: condition | Sequential Bayes Factors ($N_{min} = 4$; $N_{max} = 16$) | Repeated-measures ANOVA<br>Within-subject factor: condition<br>Random factors: participant ID, gender | <u>Inconclusive evidence</u><br>$1 < BF < 3$ = anecdotal evidence for $H_0/H_1$<br>$1 < BF < 10$ = moderate evidence for $H_0/H_1$<br><u>Conclusive evidence</u><br>$10 < BF < 30$ = strong evidence for $H_0/H_1$<br>$BF > 30$ = very strong evidence for $H_0/H_1$ |
| **Secondary Outcomes** |  |  |  |  |
| **Melatonin concentrations (S1)** | The time course ($t_1, t_2, \dots, t_n$) of melatonin concentrations on evening 1 of the protocol will differ between the three conditions. Concentrations in the constant light condition will be highest, in the yellow-bright flickering they will be lowest, the blue-dim flickering will be associated with intermediate melatonin levels.<br>Effects of interest: condition, time × condition interaction. | See sampling plan of C1 | Repeated-measures ANOVA<br>Within-subject factors: condition, time<br>Random factors: participant ID, gender | See interpretation of C1 |
| **Subjective sleepiness (S2)** | The time course ($t_1, t_2, \dots, t_n$) of subjective sleepiness ratings on the Karolinska Sleepiness Scale (KSS) will differ between the three conditions. KSS values in the constant light condition will be highest (indicating higher sleepiness), in the yellow-bright flickering they will be lowest, the blue-dim flickering will be associated with intermediate KSS<br>values.<br>Effect of interest: condition, time × condition interaction. | See sampling plan of C1 | Repeated-measures ANOVA<br>Within-subject factors: condition, time<br>Random factors: participant ID, gender | See interpretation of C1 |
| **Objective Sleepiness (S3)** | The time course (t1, t2, t3) of EEG-derived alpha (8-12 Hz)/ theta (4-7 Hz) ratio values will differ between the three conditions. Ratios in the constant light condition will be smallest (indicating higher sleepiness), in the yellow-bright flickering they will be highest, the blue-dim flickering will be associated with intermediate ratios.<br>Effects of interest: condition, time × condition interaction. | See sampling plan of C1 | Repeated-measures ANOVA<br>Within-subject factors: condition, time<br>Random factors: participant ID, gender | See interpretation of C1 |
| **Visual comfort (S4)** | The time course (t1, t2, ...tn) of visual comfort ratings will differ between the three conditions. Visual comfort ratings in the constant light condition will be highest (indicating higher comfort), in the yellow-bright flickering they will be lowest, the blue-dim flickering will be associated with intermediate visual comfort values.<br>Effect of interest: condition, time × condition interaction. | See sampling plan of C1 | Repeated-measures ANOVA<br>Within-subject factors: condition, time<br>Random factors: participant ID, gender | See interpretation of C1 |
| **PVT: Median reaction time (RT) (S5)** | The time course (t1, t2, ...tn) of median RTs will differ between the three conditions. Median RTs in the constant light condition will be highest, in the yellow-bright flickering they will be lowest, the blue-dim flickering will be associated with intermediate median RTs.<br>Effect of interest: condition, time × condition interaction. | See sampling plan of C1 | Repeated-measures ANOVA<br>Within-subject factors: condition, time<br>Random factors: participant ID, gender | See interpretation of C1 |
| **PVT: Fastest 10% RTs (S6)** | The time course (t1, t2, ...tn) of the 10% fastest RTs will differ between the three conditions. 10% fastest RTs in the constant light condition will be highest, in the yellow-bright flickering they will be lowest, the blue-dim flickering will be associated with intermediate 10% fastest RTs.<br>Effect of interest: condition, time × condition interaction. | See sampling plan of C1 | Repeated-measures ANOVA<br>Within-subject factors: condition, time<br>Random factors: participant ID, gender | See interpretation of C1 |
| **PVT: Slowest 10% RTs (S7)** | The time course (t1, t2, ...tn) of the 10% slowest RTs will differ between the three conditions. 10% slowest RTs in the constant light condition will be highest, in the yellow-bright flickering they will be lowest, the blue-dim | See sampling plan of C1 | Repeated-measures ANOVA<br>Within-subject factors: | See interpretation of C1 |
|  | flickering will be associated with intermediate 10% slowest RTs.<br>Effect of interest: condition, time × condition interaction. |  | condition, time<br>Random factors: participant ID, gender |  |
| **EEG-derived sleep onset latency (SLAT) (S8)** | The SLAT (in minutes) on evening 1 of the protocol is longer in the yellow-bright condition than in the blue-dim condition. The SLAT in both conditions are longer than in the constant light condition.<br>Effect of interest: condition | See sampling plan of C1 | Repeated-measures ANOVA<br>Within-subject factor: condition<br>Random factors: participant ID, gender | See interpretation of C1 |
| **EEG-derived slow wave activity (SWA) (S9)** | The time course (t1, t2, ...tn) of SWA (averaged power between 0.5 and 4.5 Hz at electrodes F3, F4, Fz) in each percentile of the first sleep cycle will differ between the three conditions. SWA in the constant light condition will be highest, in the yellow-bright flickering they will be lowest, the blue-dim flickering will be associated with intermediate median RTs.<br>Effect of interest: condition, time × condition interaction. | See sampling plan of C1 | Repeated-measures ANOVA<br>Within-subject factors: condition, time<br>Random factors: participant ID, gender | See interpretation of C1 |
| **Outcome-neutral measurements** |  |  |  |  |
| **Melatonin concentrations (ON1)** | The time course (t1, t2, ...tn) of melatonin concentrations on evening 1 of the protocol <u>before the light exposure</u> will *not* differ between the three conditions.<br>Effects of interest: condition, time × condition interaction. | See sampling plan of C1 | Repeated-measures ANOVA<br>Within-subject factors: condition, time<br>Random factors: participant ID, gender | See interpretation of C1 |
| **PVT: Median reaction time (RT) (ON2)** | The time course (t1, t2, ...tn) of median RTs <u>before the light exposure</u> will *not* differ between the three conditions.<br>Effects of interest: condition, time × condition interaction. | See sampling plan of C1 | Repeated-measures ANOVA<br>Within-subject factors: condition, time<br>Random factors: participant ID, gender | See interpretation of C1 |
| **Subjective sleepiness (ON3)** | The time course (t1, t2, ...tn) of subjective sleepiness ratings on the Karolinska Sleepiness Scale (KSS) <u>before the light exposure</u> will *not* differ between the three conditions. | See sampling plan of C1 | Repeated-measures ANOVA<br>Within-subject factors: | See interpretation of C1 |
|  | Effect of interest: condition, time × condition interaction. |  | condition, time<br>Random factors:<br>participant ID,<br>gender |  |
| **Brightness**<br>**(ON4)** | The time course (t1, t2, ...tn) of perceived brightness ratings <u>before the light exposure</u> will *not* differ between the three conditions.<br>Effect of interest: condition, time × condition interaction. | See sampling plan of C1 | Repeated-measures ANOVA<br>Within-subject factors:<br>condition, time<br>Random factors:<br>participant ID,<br>gender | See interpretation of C1 |

Factors: “condition” = three factor levels (constant light, blue-dim flickering, yellow-bright flickering); “time” = multiple assessments/time course within one condition (factor levels: t1, t2, ..., tn); “gender” = two factor levels (man vs. woman); “participantID” = participant code (number of factor levels depends on final sample size); BF = Bayes Factor, can be BF<sub>10</sub> or BF<sub>01</sub>. Note that we decided to implement the repeated-measures ANOVA using a linear model approach to circumvent the issue of case-wise deletion in case of missing data.

**Table 2.** Overview of the derived measure and the number of outcome measurements per participant for each hypothesis in the design table (**Table 1**).

| Measurement modality | Derived measure | Number of outcome measurements per participant |
| --- | --- | --- |
| **Primary outcome** |  |  |
| **Melatonin concentrations (C1)** | *Circadian phase shifts* | 3 (1x constant light, 1x ‘blue-dim’ flickering, 1x ‘yellow-bright’ flickering) |
| **Secondary outcomes** |  |  |
| **Melatonin concentrations (S1)** | *Time series of melatonin concentrations* | 42 (14x constant light, 14x ‘blue-dim’ flickering, 14x ‘yellow-bright’ flickering) |
| **Subjective sleepiness (S2)** | *Time series of subjective sleepiness ratings* | 42 (14x constant light, 14x ‘blue-dim’ flickering, 14x ‘yellow-bright’ flickering) |
| **Objective Sleepiness (S3)** | *Time series of objective sleepiness* | 9 (3x constant light, 3x ‘blue-dim’ flickering, 3x ‘yellow-bright’ flickering) |
| **Visual comfort (S4)** | *Time series of visual comfort ratings* | 30 (10x constant light, 10x ‘blue-dim’ flickering, 10x ‘yellow-bright’ flickering) |
| **Reaction time (S5)** | *Time series of median RT* | 42 (14x constant light, 14x ‘blue-dim’ flickering, 14x ‘yellow-bright’ flickering) |
| **(S6)** | *Time series of fastest 10% RT* | 42 (14x constant light, 14x ‘blue-dim’ flickering, 14x ‘yellow-bright’ flickering) |
| **(S7)** | *Time series of slowest 10% RT* | 42 (14x constant light, 14x ‘blue-dim’ flickering, 14x ‘yellow-bright’ flickering) |
| **Sleep EEG (S8)** | *Sleep onset latency to consistent 10 min of sleep (SLAT)* | 3 (1x constant light, 1x ‘blue-dim’ flickering, 1x ‘yellow-bright’ flickering) |
| **(S9)** | *Slow Wave Activity (SWA)* | 30 (10 percentiles ‘constant light’, 10 percentiles ‘blue-dim’, 10 percentiles ‘yellow-bright’) |

Please note that the precise number of evening measurements depends on the habitual bed time.

**Table 3.** Converging evidence for a role of S-cone-opponent circuitry.

| Study | Species | Level of Analysis | Findings |
| --- | --- | --- | --- |
| Mouland et al. (2019)<sup>24</sup> | Rodents | Behavioural | • Light modulations, which stimulate the short wavelength-sensitive S and long wavelength-sensitive L cones in an opponent fashion affect the murine circadian system differentially: Yellow-bright stimuli have stronger circadian phase-shifting effects than blue-dim stimuli.<br>• Shows that these modulations have effects independent of the melanopsin system. |
| Dacey et al. (2005)<sup>8</sup> | Primates | Anatomical | • S-cones provide an inhibitory input and the L and medium wavelength-sensitive M cones provide an excitatory input to the ipRGCs. |
| Patterson et al. (2020)<sup>9</sup> | Primates | Anatomical | • The circuit described by Dacey and colleagues is due to a specific amacrine cell. |
| Krauskopf et al. (1982)<sup>18</sup><br>Webster et al. (1994)<sup>92</sup> | Humans | Psychophysical | • Colour vision is organised by so-called ‘cardinal directions’, one of which is the S–[L+M] direction pitting S-cones against luminance (L+M) |
| Spitschan et al. (2014)<sup>10</sup><br>Cao et al. (2015)<sup>93</sup><br>Woelders et al. (2018)<sup>11</sup> | Humans | Physiological | • The pupil light response, which is fed by the ipRGCs receives an S-cone opponent input.<br>• This also confirms the circuit described by Dacey and colleagues. |
| Figueiro et al. (2004)<sup>19</sup> | Human | Neuroendocrine | • Both light sources, one with a peak at 460 nm (blue LED) and one without (mercury vapor, yellowish light), elicit melatonin suppression. The blue LED is more effective.<br>• Supports the role of S-cone-opponent signals in melatonin suppression in humans. |

Our hypothesis that flickering light would elicit a stronger effect was motivated by the psychophysical and physiological evidence^27–33^ that cones and postreceptoral mechanisms adapt under constant conditions, as well as a recent study showing that flashing light causes a stronger phase shift than continuous light under certain conditions^34^. The hypothesis that yellow-bright flickering stimuli have a stronger effect than blue-dim stimuli was based on the previously discussed animal study by Mouland and colleagues^24^. We subjected the circadian phase shift to a Bayes Factor test in a within-subjects ANOVA design and used the Sequential Bayes Factor^35^ estimation procedure to assess the evidence for the full model (incorporating the light condition type) over a null model (not incorporating the light condition type). We had planned to continuously recruit participants until the stopping criterion, or resource limit would be reached. The stopping criterion was a Bayes Factor (i.e., BF_10_ or BF_01_) of ≥10, which corresponds to the full model (i.e., light condition making a difference) being 10x more likely than the null model or vice versa. Our financially driven resource limit was *n*=16 participants. Assuming a large effect size (ES; Cohen’s *d* = 0.8) and that H1 better predicts the data than H0, simulations^36^ revealed that 62% of the simulations showed evidence for H1 with 38% being inconclusive (medium ES: 22.2% vs. 77.7%; small ES: 0.5% vs. 96.4% and 3.1% showing evidence for H0). However, we argue that, if the S– (L+M) opponent system indeed exerts a very strong effect on the circadian system, then this should be visible on a single-subject level already. In fact, previous research suggests that the assumptions about effect sizes (i.e., *d* = 0.8) for the circadian phase shifts used in the simulations outlined above may be rather conservative^37^. Thus, if the results remain inconclusive even after 16 participants, we argue that if there is an effect we have missed, such an effect is not physiologically meaningful in the average healthy individual. For the field, this is of great importance, as it would directly inform that any influence of S–(L+M) stimulation on the circadian system is very small, or subject to very strong interindividual differences. In those cases, the message is that melanopsin is the main driver of circadian photoentrainment.

### Secondary, ring-fenced hypotheses

In addition to our primary outcome, we also examined a set of secondary endpoints (see Tables **1** and **2** for a description of the outcome variables and associated hypotheses). The time series of melatonin concentrations in the evening of the first night was subjected to a repeated-measures ANOVA and the evidence for a difference in the acute melatonin-suppressive effect of light was examined using Bayes Factors (***S1***). We hypothesized that the yellow-bright stimuli would have a larger melatonin-attenuating effect than the blue-dim stimuli and that both would have a larger effect than constant light. Psychometrically, we collected sleepiness ratings using the Karolinska Sleepiness Scale^29^ (***S2***) and visual comfort ratings using a custom 9-item rating scale (***S4***), which we likewise planned to subject to a Bayesian repeated-measures ANOVA to examine differences. Objective sleepiness was assessed by ratio of absolute theta (4-7 Hz) and alpha (8-12 Hz) power at parietal and occipital electrodes during resting-state electroencephalography (EEG) with both eyes open^38^ (***S3***). We hypothesized that sleepiness (ratings) would be lower and visual discomfort ratings would be higher in the yellow-bright stimuli than in the blue-dim stimuli. Constant light was expected to be associated with highest sleepiness (ratings) and highest visual comfort ratings. We also examined changes in performance on a reaction time test, a modified auditory Psychomotor Vigilance Test (PVT)^25, 26, 39–41^, to test for differences in vigilant attention between the three lighting conditions using a Bayesian repeated-measures ANOVA. We hypothesized that yellow-bright stimuli would produce faster reaction times as measured with the PVT than blue-dim stimuli and that constant light would be associated with slowest reaction times (median, 10% fastest, 10% slowest; ***S5***, ***S6*** and ***S7***). Finally, we examined properties of sleep using polysomnography (PSG) in the night following light exposure. We specifically examined sleep onset latency to consistent 10 minutes of sleep^37, 42^ (***S8***), which we hypothesized to be longer in the yellow-bright than in the blue-dim condition and shortest in the constant light condition. We also studied slow-wave activity (SWA; delta power density between 0.5 and 4.5 Hz, ***S9***) during the first sleep cycle^25^, which we hypothesized to be decreased in the yellow-bright condition compared to the blue-dim condition and highest in the constant light condition.

## Methods

### Participants

Healthy, young male and female participants (age range 18-35 years; age range restricted to minimise age-related inter-observer variability in pre-receptoral filtering^43, 44^) were recruited, with the goal of including an equal number of men and women in the study. Inclusion criteria were a Body Mass Index (BMI) between 18.5 and 24.9 (i.e. normal weight according to WHO criteria, calculated from self-reported height and weight), an informed consent as documented by signature of the participant as well as approval of study participation by the study physician. Participants were excluded if they were pregnant (self-report), suffered from chronic or debilitating medical conditions (physical examination by the study physician), used medications impacting on visual, neuroendocrine, sleep, and circadian physiology (determined by study physician), used drugs (verified with a urine multi-drug screen during each experimental night; nal von minden GmbH, Moers, Germany), had performed shift work <3 months prior to beginning of the study, travelled across more than two time zones <1 month prior to beginning of the study, were characterised by an extreme chronotype according to the Munich Chronotype Questionnaire (MCTQ)^45^ (exclude values ≤2 or ≥7, include >2 and <7), had extremely short or long sleep duration (subjective sleep duration on workdays outside 6-10 hours according to the MCTQ), had a sleep efficiency <70% during the adaptation night (calculated as SEFF = total sleep time / time in bed using visual scoring), any indicators of a sleep disorder (self-report or during the adaptation night), or photosensitive epilepsy. Furthermore, they were excluded in case of previous enrolment in this study or if they were the investigators’ family members, employees or other dependent persons. They were also excluded if they were unable to understand and/or follow the study procedures or in case of non-compliance to sleep-wake times during the ambulatory part. Participants were furthermore screened for normal colour vision using the Cambridge Colour Test^46^ (trivector version) implemented using an iMac-based Metropsis system (Cambridge Research Systems, Rochester, UK) and were excluded if they did not have normal colour vision. Participants whose bedtime or wake time deviated by more than 30 minutes more than twice during the five days preceding each experimental visit were not empanelled in the study or excluded. All exclusion criteria are listed in Table **4**. Any exclusions were counted and reported. In case participants had to be excluded due to non-adherence to sleep-wake times during the circadian stabilisation period, but had already completed >1 study visit, none of their data was be used. Two participants were planned to be tested at the same time (for more information cf. section “Deviations from Protocol”). Participants received a remuneration of CHF1000 after successful completion of the study protocol.

**Table 4.** Systematic overview of exclusion criteria.

| **Aspect** | **Modality** | **Exclusion criterion** | **Time of assessment** |
| --- | --- | --- | --- |
| *Ability to understand study materials* | n/a | Inability to understand study materials | Pre-screening |
| *Previous exposure to study* | n/a | Prior participation in study | Pre-screening |
| *Relationship with study team* | n/a | Family members, employees or other dependent persons | Pre-screening |
| *Normal body weight* | BMI calculated from self-reported height and weight | <18.5 or >24.9 | Pre-screening |
| *Pregnancy* | Self-report | Pregnancy | Pre-screening |
| *Shift work* | Self-report | Shift work <3 months prior to Visit 1 | Pre-screening |
| *Travel across time zones* | Self-report | Travel across two time zones <1 month prior to Visit 1 | Pre-screening |
| *Chronotype* | MCTQ | $\leq 2$ or $\geq 7$ | Pre-screening |
| *Habitual sleep duration* | MCTQ | <6 or >10 hours | Pre-screening |
| *Physical health* | Physical examination by study physician |  | Adaptation night (Visit 1) |
| *Medications impacting on visual, neuroendocrine, sleep, and circadian physiology* | Physical examination by study physician |  | Adaptation night (Visit 1) |
| *Sleep efficiency* | PSG / Laboratory log | <70% (total sleep time / time in bed) | Adaptation night (Visit 1) |
| *Normal sleep* | Self-report | Any indicators of sleep disorder | Adaptation night (Visit 1) |
| *Photosensitive epilepsy* | Self-report | Self-reported photosensitive epilepsy | Adaptation night (Visit 1) |
| *Colour vision* | Cambridge Colour Test | Normal colour vision | Adaptation night (Visit 1) |
| *Core body temperature* | Core body temperature measurements using ingestible pill | Drop less than 0.2 between arrival in the laboratory and habitual bed time | Adaptation night (Visit 1) |
| *Circadian stabilisation* | Actigraphy and sleep diary | Deviation of target bed or wake time >30 minutes 2x during 5 days prior to study visit | Prior to experimental visits (Visit 2, 3 and 4) |

### Protocol

In this within-subject protocol, all participants underwent the same study procedures, which started with an initial 7-day circadian stabilisation period in which they were instructed to go to bed within 1 hour (±30 minutes) of a target bedtime and rise within 1 hour (±30 minutes) of a target wake-up time with a target sleep duration of 8 hours. Both target bedtimes and wake-up times were agreed upon with the participants to match their habitual bedtime and ensure compatibility with daily life. Compliance to the individual sleep-wake schedules was ensured using wrist actimetry (Centre Point Insight Watch; ActiGraph LLC, Pensacola, FL, USA) and sleep logs^47^. They also had to adhere to this stabilisation protocol during the ambulatory phases between further visits to the laboratory. In total, participants entered the laboratory four times. The first laboratory visit was an adaptation night, and the second, third, and fourth laboratory visits were experimental visits both including two evenings, one night, and one day (32 hours [was actually 32.5 hours; cf. “Deviations from Protocol”]) in the lab (Figure **2**).

During the first night of each of the three experimental visits, participants were exposed to the constant light (i.e., “background” condition), a yellow-bright flickering condition, or a blue-dim flickering condition between 30 minutes and 90 minutes after habitual bedtime (1-hour exposure duration). While the first visit was always the constant light condition (“background”), the order of the other two conditions was counterbalanced among participants and within each gender subgroup. Note that in the Stage 1 protocol it was incorrectly stated that the order of all three conditions would be randomised, which deviated from the information provided in Figure 2, its caption, and the hypotheses. Compliance with instructions to keep eyes open during the light exposure was checked informally through electrooculography, which had to show blinks. Experimental visits were spaced by a wash-out phase of one week (i.e., 5 nights).

On each experimental visit, participants entered the laboratory 6.5 hours prior to their habitual bedtime. Upon arrival, a small dinner was served and the EEG, electrooculogram (EOG), and electromyogram (EMG) for later sleep assessment (i.e., PSG, see below) was be placed. From 5 hours prior to habitual bedtime (HBT) until 1 h 30 min after HBT, participants provided saliva samples every thirty minutes, using Salivettes (Sarstedt), which were centrifuged at 3000 rpm for 3 minutes and frozen at –20° (was actually -28°C, cf. “Deviations from Protocol”) for later assaying.

Participants rated subjective sleepiness on the Karolinska Sleepiness Scale (KSS)^48^, completed a modified auditory Psychomotor Vigilance Test (PVT, 6 minutes; was actually 10 minutes [cf. “Deviations from Protocol”]) to assess behavioural vigilance, and rated their visual comfort with every melatonin sample. We used the German version of the Karolinska Sleepiness Scale (“Bitte bewerten Sie Ihre Müdigkeit”: “sehr wach” [1],“wach” [3], “weder wach noch müde” [5], “müde, aber keine Probleme, wach zu bleiben” [7], “sehr müde, große Probleme, wach zu bleiben, mit dem Schlaf kämpfend” [9]). Furthermore, we recorded a 3-min resting-state EEG (open eyes) just before the start of the light exposure, 30 min into light exposure, and immediately after light exposure (i.e., 27 min, 1 h, and 1 h 27 min after HBT). Following a visit to the bathroom (approx. 15 min), they went to bed 2 h after their habitual bedtime for a 6-h sleep opportunity. Following wake-up they provided another five saliva samples during the first 2 h of wakefulness. They then spent the day in the lab under controlled lighting conditions (<8 lux; fluorescent lighting) and we had planned to serve small meals every 2.5 hours (5 snacks with 0.2 of the total basic metabolic rate estimated with the Mifflin-St. Jeor equation^49^; was actually 6 snacks, cf. section “Deviations from Protocol”). In the second evening, we again started sampling melatonin in 30 min intervals 5 hours prior to HBT and obtained melatonin samples, KSS ratings, and PVT measurements until 1 h 30 min after HBT. For an illustration of the study protocol, please see Figure 2. Please note that there were no hypotheses for the KSS or PVT in the second evening, but the protocol was the same on both evenings so the KSS values are comparable. They then went bed for a 9-h recovery sleep opportunity. The protocol formally ended 1 h 30 min after HBT in the second night, however participants were offered to sleep in the lab. Throughout light exposure, participant’s compliance was be monitored using infrared cameras pointed at the participants.

Throughout the repeated laboratory visits, participants had no knowledge of external time and were not allowed to use their phones (except in emergencies) or laptops. They were allowed to read magazines, books or other material (e-readers were allowed if the corneal illuminance was <8 lux). Small gaming devices such as GameBoys (monochrome display), or table-top and card games were be provided. Participants were allowed to listen to podcasts provided they had a device that did not tell the time.

#### Adaptation night

Participants came for an initial adaptation night (8-h sleep opportunity) to the laboratory to screen for potential sleep disorders (see below for the PSG setup) and to make them acquainted with the laboratory setting. Participants showing signs of sleep disorders or presenting with sleep efficiency <70% would have been excluded from the study.

#### Polysomnography

For resting-state and nocturnal PSG recordings (i.e., EEG, EOG, and EMG) we used ambulatory BrainProducts LiveAmp® devices. We planned to record PSG at a sampling rate of 250 Hz (was actually 500 Hz; cf. “Deviations from Protocol”) from 21 scalp channels mounted in a cap (EasyCap®) and 4 EOG channels with FCz as the online reference. In addition, we placed two mastoid electrodes for later re-referencing according to the sleep staging criteria of the American Association of Sleep medicine, as well as two chin EMG electrodes and two electrocardiogram (ECG) electrodes^50^. During the adaptation nights, we additionally recorded from two EMG electrodes on the tibialis anterior muscle of one leg to screen for nocturnal leg movements, a respiration belt, and a nasal cannula to screen for respiratory problems.

#### Visual comfort questionnaire

Participants were planned to be asked to rate or respond to various aspects of the light exposure using a 6-question, 7-item Likert scale questionnaire^12^. This questionnaire was administered in German. The questions were planned to be about the comfort of light (“Allgemein ist das Licht angenehm”; überhaupt nicht [1] – sehr stark [7]), the perceived brightness (“Wie empfinden Sie die Helligkeit des Lichtes?”; sehr dunkel [1] – sehr hell [7]), light level preference (“Ich hätte es lieber …”; deutlich dunkler [1] – deutlich heller [7]), glare (“Dieses Licht blendet mich”; überhaupt nicht [1] –sehr stark [7]), the perceived colour temperature (“Wie empfinden Sie die Lichtfarbe?”; sehr kalt [1] – sehr warm [7]) and general well-being (“Wie fühlen Sie sich im Moment?”; unwohl [1] – wohl [7]). For deviations from the planned questionnaire, please see the “Deviations from Protocol” section.

### Light exposure and stimuli

#### Evening light stimuli

Light exposure was planned to be delivered from a vertical front lighting panel (width 220 cm, height 140 cm) that consists of 24 LED panels (RGBW), each containing 144 LEDs (i.e. total of 3456 LEDs) covered by diffusing material^51, 52^. Two participants were planned to sit next to each other at a distance of 1.5 m from the wall, facing the wall. The LED primaries have the following properties (peak wavelength±FWHM; CIE 1931 *xy* chromaticity): blue (462±23 nm; 0.14, 0.06), green (518±32 nm; 0.16, 0.72), red (631±16 nm; 0.70, 0.30), white (peaks 446 nm and 560 nm; 0.33, 0.34). Due to the Coronavirus pandemic, an alternative setup with two separate laboratories had to be used. For details, please see the “Deviations from Protocol” section.

The wall was planned to be calibrated from a vantage point centred between the two participants, and both irradiance and radiance measurements were planned to be taken. To characterise spectral shifts of the LED wall, we planned to measure the spectrum of each of the RGBW primaries at 16 settings at 8-bit resolution.

Stimuli were either a constant light matched in its photoreceptor activation profile to that of daylight at 6500K (D65, used during the first experimental visit; “background condition”; Figure **1*f*** [note that the stage 1 protocol wrongly stated that this light was used during the adaptation night and the second night of each experimental visit]), or sinusoidally flickering light stimulating the S-cones in an balanced but opponent fashion with luminance (blue-dim: planned +37% S-cone contrast, planned –37% L+M contrast from background, Figure **1*g***; yellow-bright: planned –37% S-cone contrast; planned +37% L+M cone contrast from background, Figure **1*h***), alternating flickering for 30 seconds to constant background for 30 seconds. The flicker frequency was 1 Hz so as to provide a continuous tonic signal to the cones which might otherwise adapt to a continuous light biased towards either –S+(L+M) or +S–(L+M) (Figure **1*e***). This frequency was also used in the habituation stimulus in a seminal study determining the cardinal directions of colour space^18^. To avoid adaptation to flicker, we flickered for 30 seconds, then held the light constant at background for 30 seconds, corresponding to the approximate time scale of habituation in the purported S-opponent psychophysical channel^18^. All stimulus conditions are summarized in terms of their nominal photoreceptor excitation^53^ in Table **5**.

**Table 5.** Stimulus characteristics. Overview of the irradiance-derived α-opic responses (in mW/m2) for the three experimental screen light conditions with ambient dim light. Photopic illuminance was 93.5 lux (background), 123.5 lux (yellow-bright), and 67.1 lux (blue-dim). Radiance-derived chromaticity values (CIE 1931 xy standard observer for a 2° field) were x = 0.25 and y = 0.3 for the background, x = 0.26 and y = 0.4 for the yellow-bright, and x = 0.23 and y = 0.2 for the blue-dim condition. Measures were taken at a distance of 68 cm from the screen at a height of 118 cm, that is, from the observer’s point of view. Values were calculated using the luox app^94^. For a reference, we also report values for daylight (D65) at 93.5 lux.

| | α-opic irradiances [mW/m<sup>2</sup>] | | | | CIE 1931 xyY | | |
| --- | --- | --- | --- | --- | --- | --- | --- |
|  | L cones | M cone | S-cone | Melanopsin | x | y | Illuminance [lx] |
| **Background** | 159.22 | 160.83 | 84.90 | 212.91 | 0.25 | 0.30 | 93.53 |
| **+S-(L+M)** | 123.67 | 124.21 | 110.63 | 219.60 | 0.23 | 0.20 | 67.01 |
| *Contrast* | -22.33% | -22.77% | 30.31% | 3.15% |  |  |  |
| **-S+(L+M)** | 200.71 | 204.06 | 63.21 | 216.83 | 0.26 | 0.40 | 123.49 |
| *Contrast* | 26.06% | 26.88% | -25.54% | 1.84% |  |  |  |
| Daylight (D65) | 162.89 | 145.57 | 81.71 | 132.60 | 0.31 | 0.33 | 100 |

**Table 6.** Overview of the irradiance-derived equivalent daylight (D65) illuminances (in lux) for the three experimental screen light conditions with ambient dim light. Measures were taken at a distance of 68 cm from the screen at a height of 118 cm, that is, from the observer’s point of view. Values were calculated using the luox app^94^.

|  | Equivalent Daylight Illuminance [EDI; lux] |  |  |  |
| --- | --- | --- | --- | --- |
|  | L cones | M cone | S-cone | Melanopsin |
| **Background** | 97.75 | 110.74 | 103.88 | 160.54 |
| **+S-(L+M)** | 75.92 | 85.32 | 135.36 | 165.59 |
| *Contrast* | -22.33% | -22.95% | 30.30% | 3.05% |
| **-S+(L+M)** | 123.22 | 140.17 | 77.34 | 163.50 |
| *Contrast* | 26.06% | 26.58% | -25.55% | 1.81% |

**Table 7.** Overview of the radiance-derived α-opic responses (in mW/m^2^*sr) for the three experimental screen light conditions with ambient dim light. Photopic luminance was 298.92 cd/m^2^ (background), 404.37 cd/m^2^ (yellow-bright), and 202.32 cd/m^2^ (blue-dim). Radiance-derived chromaticity values (CIE 1931 xy standard observer for a 2° field) were x = 0.25 and y = 0.3 for the background, x = 0.26 and y = 0.4 for the yellow-bright, and x = 0.23 and y = 0.2 for the blue-dim condition. Measures were taken at a distance of 68 cm from the screen at a height of 118 cm, that is, from the observer’s point of view. Values were calculated using the luox app^94^.

| | α-opic radiances [mW/m<sup>2</sup>*sr] | | | | CIE 1931 xyY | | |
| --- | --- | --- | --- | --- | --- | --- | --- |
|  | L cones | M cone | S-cone | Melanopsin | x | y | Luminance [cd/m<sup>2</sup>] |
| **Background** | 508.24 | 543.67 | 304.10 | 755.63 | 0.25 | 0.30 | 298.92 |
| +S-(L+M) | 377.36 | 410.45 | 394.49 | 775.19 | 0.23 | 0.20 | 202.32 |
| *Contrast* | -25.75% | -24.50% | 29.72% | 2.52% |  |  |  |
| -S+(L+M) | 653.75 | 696.25 | 224.66 | 767.68 | 0.26 | 0.40 | 404.37 |
| *Contrast* | 28.63% | 28.06% | -26.12% | 1.59% |  |  |  |

To find the settings on the RGBW channels of the lighting panel, we implemented an optimisation minimising the squared error between desired and actual contrasts using ‘fmincon’ routine as implemented in MATLAB (Mathworks®, Natick, USA). Critically, our stimuli were designed to produce no differential stimulation of melanopsin-expressing intrinsically photosensitive retinal ganglion cells (0%).

#### Light during the scheduled day

During the scheduled day (i.e., from wake-up until the end of the protocol), participants were planned to be in dim light (<8 lux) provided by fluorescent lighting.

### Data collection & processing

All data handling was done in R^54^, except for EEG data. EEG data processing was done in MATLAB (The Mathworks, Natick, MA, USA) using the ‘Fieldtrip’ toolbox for MATLAB^55^ first before the results were exported for statistical analyses in R.

#### Hormone concentrations (melatonin)

Analyses of salivary melatonin were planned to be done with enzyme-linked immunoabsorbent assays (ELISAs) in an in-house laboratory. We planned to use ELISA kits (NovoLytiX, Pfeffingen l, Switzerland), which have a minimal detection limit (limit of quantification) of at least 1.6 pg/mL. Eventually, a radioimmunoassay (RIA) was used instead of an ELISA, cf. “Deviations from Protocol” section. The dim light melatonin onset (DLMO) was determined by fitting evening melatonin profiles by a piecewise linear-parabolic function using the hockey-stick algorithm (planned: v2.4, used version: 2.5; cf. “Deviations from Protocol”) to calculate the DLMO^56^.

#### Control variables

For the PVT data, trials with reaction times ≥100 ms were considered valid trials^39^. We then computed the median reaction times as well as reaction times for the fastest and slowest deciles of valid trials. KSS ratings were analysed without further pre-processing steps. As for the neuroendocrine responses, statistical evaluation was planned to be with repeated-measures ANOVAs and planned contrasts (note that we implemented an ANOVA-like approach using linear models to circumvent case-wise deletion; cf. “Deviations from Protocol”).

#### EEG analyses

Following data acquisition, the signal from EEG channels was filtered and re-referenced to a linked mastoids reference. For the analyses of objective sleepiness, we first corrected for eye movements using an independent component analysis (ICA), which was followed by manual exclusion of other artefacts (e.g. muscle artefacts). Subsequently, artefact-free data was segmented into 2-second time bins and subjected to Fast-Fourier transformations (FFT) using the ‘mtmfft’ function as implemented in the Fieldtrip® toolbox yielding a frequency resolution of 0.5 Hz. We then extracted absolute power in the theta (4-7 Hz) and alpha (8-12 Hz) frequency range and averaged across parietal and occipital electrodes (i.e., P3, Pz, P4, O1, Oz, O2) and segments within each frequency band. We then computed the alpha/theta ratio. For sleep analyses, we had planned to score sleep semi-automatically with an algorithm (The SIESTA Group, Vienna, Austria; ^57, 58^). These analyses were planned to be based on EEG data from electrodes F3, F4, C3, C4, O1, and O2 (down sampled to 128 Hz and re-referenced to the contralateral mastoid electrode) along with the signal from the EOG channels that have been placed according to Rechtschaffen and Kales (RK) criteria and the chin EMG signal. However, for budgetary reasons, we eventually used the Somnolyzer algorithm as implemented in the Philips Respironics Sleepware G3 software instead of the SIESTA scoring (cf. section “Deviations from Protocol”). Additionally, we computed EEG slow-wave activity (SWA) (i.e., delta power density between 0.5 and 4.5 Hz) as an indicator of sleep propensity across the night within each NREM part of a sleep cycle. To this end, we computed EEG SWA for each decile of the NREM part of the first NREM-REM cycle^59, 60^. For the computation of delta power density, artefact-free data was segmented into 2-second time bins and subjected to FFTs yielding a frequency resolution of 0.5 Hz. Thereafter, FFT results were averaged in the 0.5-4.5 Hz range at frontal electrodes F3, Fz, and F4 within each percentile of each NREM cycle. For each NREM cycle, the analyses thus yielded 10 measures per participant. The analysis procedure described here is based on a publication by Chellappa and colleagues^25^.

### Statistical data analysis

All statistical analyses were performed in R^54^. Statistical tests were performed the *BayesFactor* package^61, 62^, and we planned to implement an ANOVA design (function ‘anovaBF’). Note that we implemented an ANOVA-like approach using linear models instead of the classic ANOVA function to circumvent case-wise deletion in case of missing data, for details see the section “Deviations from Protocol”. The analytic strategy is described in Table **1**. Participant ID and gender were entered as random factors.

#### Exclusion criterion

Only data from participants for whom the light exposure took place during the rising arm of the melatonin curve and after the DLMO were included in the analyses.

#### Bayesian sampling strategy using Sequential Bayes Factors

Our sample size and recruitment approach were intimately linked to the analytic strategy. We intended to collect data until obtaining sufficient evidential strength for our primary hypothesis (C1) that the circadian phase shift is larger in the yellow-bright flickering condition than in the blue-dim flickering condition and the constant light condition, or until reaching our resource limit (*n*=16). Formally, this would have been achieved by calculating Sequential Bayes Factors^35^. First, the repeated-measures ANOVA described above would have been run sequentially after data from each new participant was included in the data set. Participants would then have continuously been recruited for the study until sufficient evidence for the full model would have been reached (male, female alternating). We consider a BF of 10 as “strong” evidence, following standard categorisations: 1 < BF < 3 –anecdotal evidence, 3 < BF < 10 – moderate, 10 < BF < 30 –BF > 30 – very strong evidence. According to the Sequential Bayes Factor approach, if this threshold was reached for the primary hypothesis, the study would have been halted, and the evidence strength would have been reported according to the Bayes Factor. Otherwise, the study would have been halted if the resource limit is reached (*n*=16). A minimum number of four participants (two men, two women) were planned to participate in the study, in case the evidence threshold is reached with one, two or three participants already.

### Deviations from Protocol

#### Protocol

The protocol lasted for 32.5 hours instead of the 32 hours previously described in the text, because we initially did not take into account that the last melatonin sample and round of assessments would take approx. 15-20 minutes.

Saliva samples were frozen at -28°C rather than the previously stated -20°C, because we used a different freezer.

Instead of a 6-min version of the psychomotor vigilance task, we used a 10-min version. Specifically, the 10-min version has been associated with increased sensitivity to modulations of both sleep homeostatic and circadian drives and to improvements in alertness after wake-promoting interventions^39^. A shorter version has only been recommended if the protocol does not allow for the 10-min version^63, 64^. As the protocol allowed us to use the longer version, we felt using the longer version would improve the validity of our measurements.

Instead of five snacks every 2.5 hours with 0.2 of the total basic metabolic rate estimated with the Mifflin-St. Jeor equation^49^, we served six snacks of 0.17 of the total basic metabolic rate every 2.5 hours. This was to shorten the time between the last snack and the bedtime to 5.5 instead of 8 hours, and thus to prevent participants from becoming too hungry in the evening. Additionally, we allowed participants to use their phones during a “social hour” scheduled four hours after wake-up as we have previously experienced this to increase adherence to long study protocols. Importantly, participants wore orange-tinted blue-blocking glasses during this time and were instructed to keep display brightness as low as possible.

#### Polysomnography

We recorded data at a sampling rate of 500 Hz instead of the previously planned 250 Hz with 500 Hz being the standard sampling rate in our laboratory. As we were only interested in frequencies up to 12 Hz, this change has however not affected the analyses or results.

#### Visual comfort questionnaire

Due to internal communication issues, the visual comfort questionnaire slightly deviated from the planned version and corresponded to a version used in an earlier study conducted in the laboratory. In more detail, instead of a 6-item Likert scale, a 5-item Likert scale was used. Furthermore, the questions slightly deviated from the initially planned ones. Importantly though, the relevant questions to assess visual comfort were still included. Contrary to what had been planned, participants were not asked to rate their light level preference (“I would prefer it to be lighter/darker”) and their general wellbeing (“I feel well/unwell”). Instead, participants rated the light according to how wake-promoting vs. tiring it was (“Die Beleuchtung in diesem Zimmer…”; hilft mir, wach zu bleiben [1] – ermüdet mich stark [5]; Engl.: “The lighting in this room…”; helps me to stay awake [1] – is very tiring [5]) and whether they felt it helped them to concentrate (“Die Beleuchtung in diesem Zimmer…”; hilft mir, mich besser zu konzentrieren [1] – stört meine Konzentrationsfähigkeit [5]; Engl.: “The lighting in this room…”; helps me to concentrate [1] – disturbs my ability to concentrate [5]). These two questions were however not (planned to be) included in the calculation of the visual comfort score, the relevant questions were still asked.

#### Light exposure and stimuli

Due to the Coronavirus pandemic, it was not feasible to have two participants sit in front of the planned LED wall for several hours next to each other. Thus, we used two custom-made displays, which provided a stimulus that was functionally equivalent to the originally planned stimulus, in two separate comparable (sleep) laboratories. Specifically, light exposure delivered through a modified 27-inch display consisting of a total of five sets of primaries, with peak spectral emissions at 430, 480, 500, 550 and 630 nm^65, 66^. The backlights were independently controlled at 8-bit resolution using DMX. For an illustration of the laboratory setup, please see Photo 1.

Due to some technical differences in the device primaries between the planned LED wall and the displays, the validated contrast between the “blue-dim” and the “yellow-bright” conditions was ±25% instead of the originally planned nominal contrast of ±37% (blue-dim: +25% S-cone contrast, –25% L+M contrast from background, Figure **1*g***; yellow-bright: – 25% S-cone contrast; +25% L+M cone contrast from background, Figure **1*h***). This validated contrast (±25%), although lower, however still yields a psychophysically substantial and physiologically meaningful contrast on the postreceptoral channel.

#### Light during the scheduled day

Sitting accommodations and furniture were placed in a way to prevent illuminances higher than 8 lux and participants were instructed and, if necessary, to not take positions in the room associated with higher illuminance levels. Nevertheless, for short periods of time, illuminances may have exceeded 8 lux (cf. Table **9**).

#### Hormone concentrations (melatonin)

Analyses of salivary melatonin were done with radioimmunoassay (RIA) rather than ELISAs in the NovoLytiX laboratory rather than an in-house laboratory. RIA and ELISAs by NovoLytiX generally reveal comparable results, but the laboratory uses RIA as analyses are less time-consuming. The RIA has an analytical sensitivity of 0.2 pg/mL and a limit of quantification of 0.9 pg/mL. For the calculation of the DLMO, we used version 2.5 of the hockey-stick algorithm (planned: v2.4), which was the most recent one available at the time of the data analysis^56^.

#### EEG analyses

Unlike originally planned we did not use the service provided by the SIESTA group for budgetary reasons. Instead, sleep was scored automatically with the Somnolyzer algorithm as implemented in the Philips Respironics Sleepware G3 software v. 4.0.1.0; ^57, 58^. Importantly, the Somnolyzer algorithm is based on the algorithm used by the SIESTA group and an automatic scoring software that is FDA cleared. Due to the change of the algorithm, analyses also included data from electrodes P3 and P4 in addition to F3, F4, C3, C4, O1, and O2. Furthermore, the data did not have to be downsampled to 128 Hz prior to analysis. However, the G3 software required us to apply some filtering prior to the sleep stage analyses. Filtering was according to AASM criteria^67^ with the EEG and EOG signals being filtered between 0.3 and 35 Hz and the EMG between 10 and 100 Hz with a Notch filter at 50 Hz to remove line noise.

#### Statistical data analyses

Instead of a classic ANOVA design using the function ‘anovaBF’, we implemented an ANOVA-like design using linear models and the function ‘lmBF’. We chose to use linear models instead of the classic ANOVA function as this would have resulted in a case-wise deletion in case of missing data (e.g., if data from even just one out of 14 PVTs per evening were not available or one DLMO out of the 3 conditions could not be determined).

## Results

To characterise spectral shifts of the display with different driving input, we measured the spectrum of each of the primaries using a Jeti spectraval 1501 (JETI Technische Instrumente GmbH, Jena, Germany). For an overview of the irradiance and radiance-derived α-opic responses for the three experimental screen light conditions and the corresponding irradiance and radiance-derived equivalent daylight (D65) illuminances, melanopic EDI, as well as luminances, please see Tables **5-8**. For an overview of the irradiance-derived α-opic responses, equivalent daylight (D65) illuminances (in lux), and melanopic EDI of the light during the scheduled day, please see Tables **9** and **10**.

**Table 8.** Overview of the radiance-derived equivalent daylight (D65) luminances (in lux) for the three experimental screen light conditions with ambient dim light. Measures were taken at a distance of 68 cm from the screen at a height of 118 cm, that is, from the observer’s point of view. Values were calculated using the luox app^94^.

|  | Equivalent Daylight Luminance [EDL; cd/m<sup>2</sup>] |  |  |  |
| --- | --- | --- | --- | --- |
|  | L cones | M cone | S-cone | Melanopsin |
| **Background** | 312.01 | 373.45 | 372.09 | 569.77 |
| +S-(L+M) | 231.73 | 281.94 | 482.68 | 584.51 |
| *Contrast* | -25.73% | -24.50 | 29.72% |  |
| -S+(L+M) | 401.34 | 478.25 | 274.88 | 578.85 |
| *Contrast* | 28.63% | 28.06% | -26.13% |  |

**Table 9.** Overview of the irradiance-derived α-opic responses (E_e_; in mW/m^2^) as well as irradiance-derived chromaticity values (CIE 1931 xy standard observer for a 2° field) for the ambient light at different locations in the room, that is, the situation during the day and between assessments in the evening. Measurements were taken from the observer’s point of view. Values were calculated using the luox app^94^. For a picture of the setup, please see photo 1.

| | α-opic irradiances [mW/m<sup>2</sup>] | | | | CIE 1931 xyY | | |
| --- | --- | --- | --- | --- | --- | --- | --- |
|  | L cones | M cone | S-cone | Melanopsin | x | y | Illuminance [lx] |
| At desk with display (1) | 13.04 | 8.45 | 1.05 | 3.54 | 0.49 | 0.42 | 7.82 |
| On the sofa facing away from the ambient light (2) | 10.05 | 6.54 | 0.79 | 2.74 | 0.49 | 0.43 | 6.03 |
| At the desk facing the surface of the desk (3) | 37.99 | 25.14 | 3.49 | 11.07 | 0.48 | 0.42 | 22.85 |
| On the bed, facing the opposite wall (4) | 21.76 | 14.32 | 1.94 | 6.22 | 0.49 | 0.42 | 13.07 |

**Table 10.** Overview of the irradiance-derived equivalent daylight (D65) illuminances (in lux) for the ambient light at different locations in the room, that is, the situation during the day and between assessments in the evening. Measurements were taken from the observer’s point of view. Values were calculated using the luox app^94^. For a picture of the setup, please see photo 1.

|  | Equivalent Daylight Illuminance [EDI; lux] |  |  |  |
| --- | --- | --- | --- | --- |
|  | L cones | M cone | S-cone | Melanopsin |
| At desk with display (1) | 8.00 | 5.80 | 1.28 | 2.67 |
| On the sofa facing away from the ambient light (2) | 6.17 | 4.49 | 0.97 | 2.07 |
| At the desk facing the surface of the desk (3) | 23.32 | 17.27 | 4.28 | 8.35 |
| On the bed, facing the opposite wall (4) | 13.36 | 9.83 | 2.37 | 4.69 |

In total, 47 individuals completed the online screening, and 18 participants were invited to participate in the study. Two volunteers decided to not continue with the study after the adaptation night for personal reasons. Thus, sixteen healthy, young male and female participants (mean: 25.5±2.7 years; 8 men and 8 women) were recruited. Data acquisition took place continuously between March and December 2022 with a break of 4 weeks in August 2022 (for more details on the distribution of participants across the acquisition period incl. subjectively reported light history on the day of the experimental visit, please see the supplemental material S3 and the laboratory log). Participants always entered the lab on the same day of the week.

For all analyses, we report BF_10_, i.e., the likelihood of the data under H1 compared to H0. For the interpretation of the BFs, please see Table **11**.

**Table 11:**
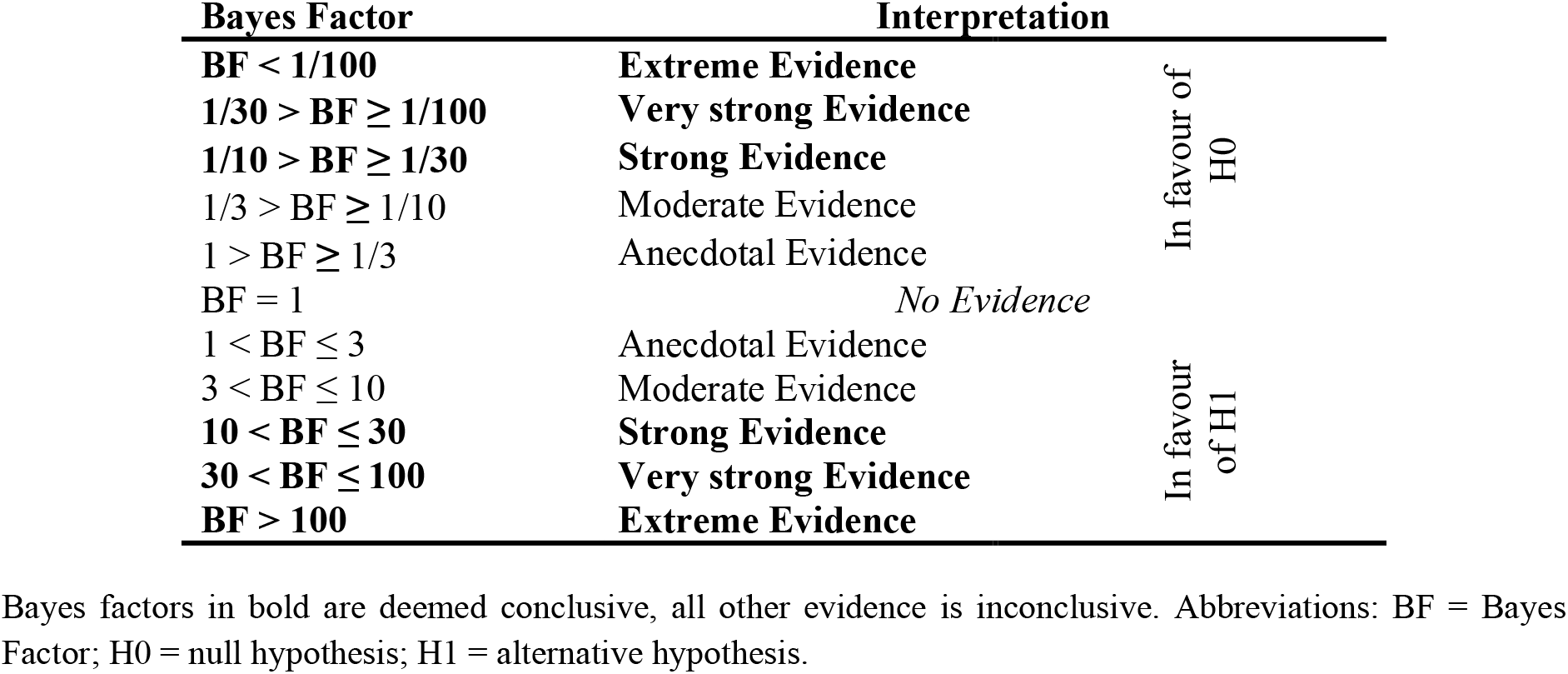
Interpretation of Bayes Factors according to Jeffreys (1961).

### Primary Outcome

#### Dim-light melatonin onset (DLMO; C1)

To determine the DLMO, we used all available evening data points. We chose a default Area of Interest Upper Border or threshold of 5 and selected “Hockeystick Time”. In rare cases, the threshold had to be adapted for some individuals. CB, CC, and MS independently performed the DLMO analyses with CC and MS being blinded to the light exposure conditions. Any divergences were resolved in a final discussion with Dr. Mirjam Münch, a colleague at the Centre for Chronobiology, who is very experienced with applying the hockey-stick algorithm. In three cases, it was not possible to determine a DLMO as no clear increase in melatonin concentrations could be identified.

Contrasting our primary hypothesis, analyses yielded moderate and thus inconclusive evidence against the hypothesis that there was a condition difference (BF_10_ = 0.3). The data were approximately 3.4 times more likely to occur under H0 than under H1. The mean phase shift was 52.0 min in the background condition (range -42.3 to 164.4 min), 41.94 min in the blue-dim condition (range -22.8 to 104.1 min), and 33.8 min in the yellow-bright condition (range -29.7 to 101.7 min). Table **12** provides an overview of the condition mean (intercept) and deviations from the intercept sampled from the posterior distribution. For a graphical illustration of the results, please see Figure **3-a**. For an illustration of the melatonin profiles on evenings 1 and 2 in each condition, see Suppl. Fig. **1**.

**Figure 3.**
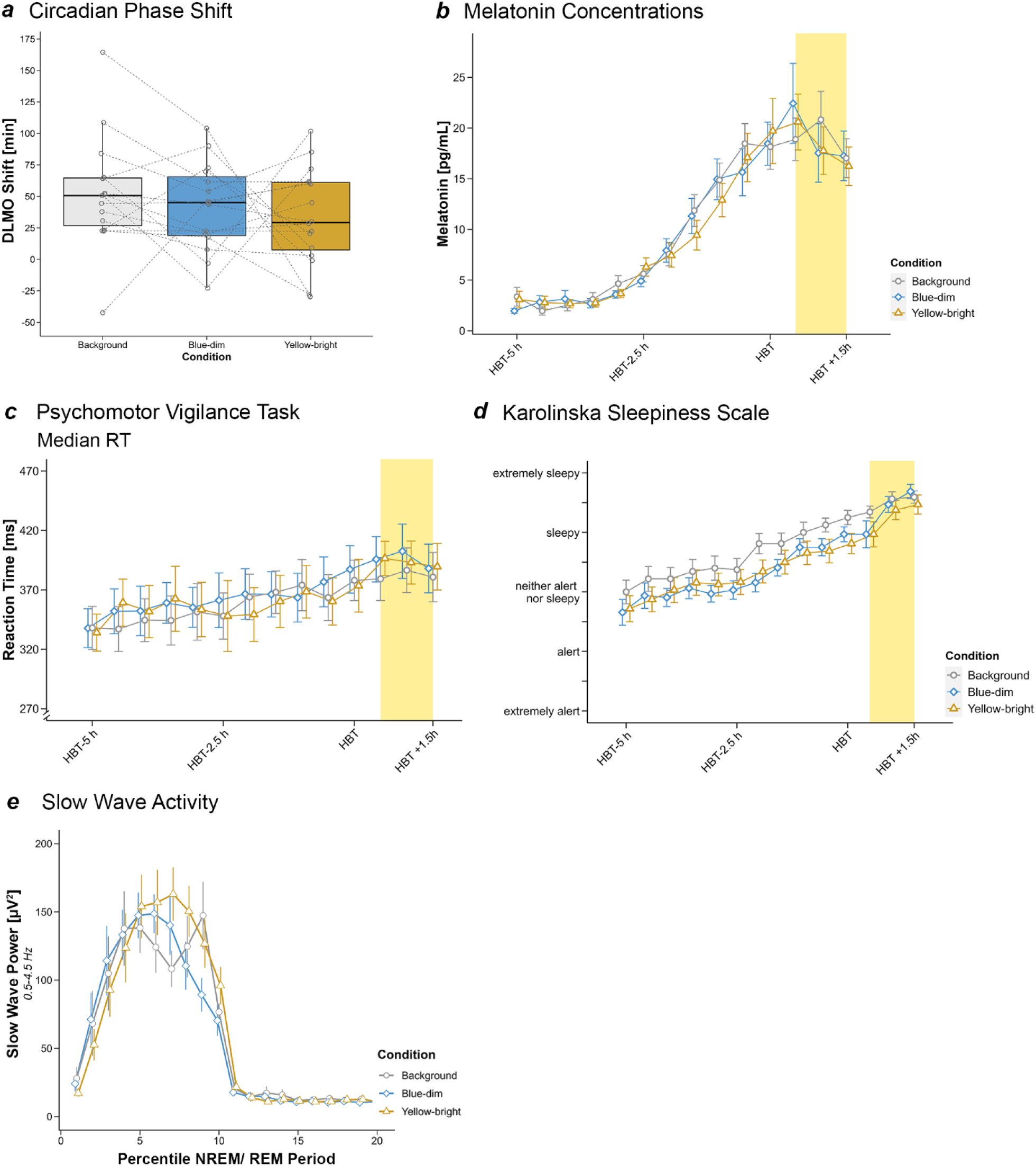
***a*** Circadian phase shift as assessed by a shift in dim light melatonin onset (DLMO) from evening 1 to evening 2 of each experimental visit. Analyses yielded inconclusive evidence against the hypothesis of a condition difference (see main text for details). The lower and upper hinges of the boxplots correspond to the 25 % and 75 % quartiles, the thick black line indicates the median. Whiskers extend to the lowest/largest value at most 1.5× the interquartile range (IQR) from the hinges. Gray circles represent individual values of participants and the dotted lines connect data points contributed by each participant. ***b*** Time course of melatonin concentrations during the first evening in the laboratory. We show the mean with error bars representing the standard error. Analyses did not yield conclusive evidence in favour of a condition difference or a time × condition interaction during the light exposure (as indicated by the yellow box) and conclusive evidence against such differences before the beginning of the light exposure. ***c*** Time course of the median reaction times on the psychomotor vigilance test (PVT) during the first evening in the laboratory. The figure shows median reaction times and 95% confidence intervals. There was conclusive evidence in favour of a condition difference before as well as during the light exposure (as indicated by the yellow box) with median reaction times being faster in the background condition (always experimental visit 1) compared to the yellow-bright and the blue-dim conditions. ***d*** Time course of subjective sleepiness ratings on the Karolinska Sleepiness Scale (KSS) during the first evening in the laboratory. The figure shows mean reaction times and error bars indicate standard errors. Evidence for a condition difference or a time × condition interaction during the light exposure (as indicated by the yellow box) remained inconclusive while there was conclusive evidence for a condition difference before the beginning of the light exposure. Specifically, participants felt more tired in the background condition corresponding to the first lab visit compared to the blue-dim or the yellow-bright conditions. ***e*** Slow-wave activity as an indicator of homeostatic sleep pressure across the first sleep cycle. The time course relates to the percentiles of the NREM period (numbers 1-10 on the x-axis) and the REM period (numbers 11-20 on the x-axis) of the first sleep cycle after sleep onset. The data points represent the mean slow wave power between 0.5 and 4.5 Hz with error bars indicating standard errors. Analyses yielded conclusive evidence against a condition difference and inconclusive evidence for a time × condition interaction.

**Table 12:** Shift in DLMO in the three light exposure conditions sampled from the posterior distribution.

| Effect | Estimate | SD | 95% CI<br>(lower; upper) |
| --- | --- | --- | --- |
| Condition mean (intercept) | 41.7 | 53.0 | -46.9; 128.0 |
| Background | 6.8 | 6.6 | -6.2; 20.6 |
| Blue-dim | -0.8 | 6.7 | -13.9; 12.2 |
| Yellow-bright | -6.0 | 6.5 | -19.6; 6.5 |

Abbreviations: SD = Standard deviation of the estimate; CI = Credible Interval. For each condition, we report differences from the intercept.

### Secondary Outcomes

#### Melatonin Concentrations (S1)

Analyses of melatonin values yielded strong evidence against a condition difference during the light exposure (BF_10_ = 0.09) suggesting that the data were approximately 11 times more likely to occur under H0. This result was largely confirmed using analyses based on rank-transformed data, which yielded moderate evidence against H1 (BF_10 rank-based_ = 0.14). For an overview of the condition mean (intercept) and deviations from the intercept sampled from the posterior distribution, please see Table **13**. There was moderate (inconclusive) evidence against a condition × time interaction (BF_10_ = 0.13; BF_10 rank-based_ = 0.13). Please see Figure **3-b** for an illustration of the results and Suppl. Fig. **1** for an illustration of the melatonin concentrations on evenings 1 and 2.

**Table 13:** Melatonin values in the three light exposure conditions sampled from the posterior distribution.

| Effect | Estimate | SD | 95% CI<br>(lower; upper) |
| --- | --- | --- | --- |
| Condition mean (intercept) | 18.7 | 8.0 | 5.0; 32.0 |
| Background | 0.15 | 0.6 | -1.0; 1.3 |
| Blue-dim | 0.3 | 0.6 | -0.9; 1.5 |
| Yellow-bright | -0.4 | 0.6 | -1.6; 0.7 |

Abbreviations: SD = Standard deviation of the estimate; CI = Credible Interval. For each condition, we report differences from the intercept.

#### Subjective Sleepiness (S2)

Analyses yielded anecdotal and thus inconclusive evidence against a condition difference regarding the subjectively reported sleepiness on the Karolinska Sleepiness Scale (KSS) during the light exposure. Under the H0, the data was approx. 1.6 times more likely than under the H1 (BF_10_ = 0.62; BF_10 rank-based_ = 0.23). For the condition mean (intercept) and deviations from the intercept sampled from the posterior distribution, please see Table **14**. Furthermore, there was moderate (inconclusive) evidence against a time × condition interaction (BF_10_ = 0.27; BF_10 rank-based_ = 0.14). For a graphical illustration, see Figure 3 D and for an illustration of the KSS values on evenings 1 and 2 in each light condition, please see Suppl. Fig. **6** (note this was not part of the analysis plan, wherefore we refrain from a statistical analysis).

**Table 14:** Subjective sleepiness ratings on the Karolinska Sleepiness Scale (KSS) during the light exposure in the three light conditions sampled from the posterior distribution.

| Effect | Estimate | SD | 95% CI<br>(lower; upper) |
| --- | --- | --- | --- |
| Condition mean (intercept) | 7.77 | 1.4 | 5.5; 10.1 |
| Background | -0.01 | 0.1 | -0.01; 0.4 |
| Blue-dim | 0.2 | 0.1 | -0.2; 0.2 |
| Yellow-bright | -0.19 | 0.1 | -0.4; 0.02 |

Abbreviations: SD = Standard deviation of the estimate; CI = Credible Interval. For each condition, we report differences from the intercept.

#### Objective Sleepiness (S3)

We assessed objective sleepiness, i.e., the alpha (8-12 Hz) to theta (4-7 Hz) ratio during a 3-min Karolinska Drowsiness Test (KDT) with eyes open at the beginning, after 30 min, and at the end of the 1-h light exposure at parieto-occipital electrodes. There was moderate (inconclusive) evidence against a condition effect with the data being approx. 3.5 times more likely under H0 than under H1 (BF_10_ = 0.28; BF_10 rank-based_ = 0.09). For an overview of the condition mean (intercept) and deviations from the intercept sampled from the posterior distribution, please see Table **15**. Analyses additionally yielded strong evidence against a time × condition interaction (BF_10_ = 0.06; BF_10 rank-based_ = 0.08).

**Table 15:** Objective sleepiness (i.e., Alpha [8-12 Hz]/ Theta [4-7 Hz] ratio) during the Karolinska Drowsiness Tests (KDT) at the beginning, 30 min into, and at the end of the light exposure in the three light conditions sampled from the posterior distribution.

| Effect | Estimate | SD | 95% CI<br>(lower; upper) |
| --- | --- | --- | --- |
| Condition mean (intercept) | 1.1 | 0.42 | 0.5; 1.8 |
| Background | -0.04 | 0.03 | -0.1; 0.01 |
| Blue-dim | 0.004 | 0.03 | -0.05; 0.05 |
| Yellow-bright | 0.03 | 0.03 | -0.01; 0.09 |

Abbreviations: SD = Standard deviation of the estimate; CI = Credible Interval. For each condition, we report differences from the intercept.

#### Visual Comfort (S4)

There was moderate (inconclusive) evidence against a difference between the conditions regarding visual comfort experienced during the light exposure (i.e., how pleasant/ bright/ glaring the light was, and how warm/cold the light colour was). The data was approximately 6 times more likely under the H0 than under the H1 (BF_10_ = 0.16). Table **16** provides an overview of the condition mean (intercept) and deviations from the intercept sampled from the posterior distribution. For the condition × time interaction, there was moderate evidence in favour of H1 (BF_10_ = 7.68). For a graphical illustration, see Suppl. Fig. **2**.

**Table 16:** Visual comfort scores during the light exposure in the three light conditions sampled from the posterior distribution.

| Effect | Estimate | SD | 95% CI<br>(lower; upper) |
| --- | --- | --- | --- |
| Condition mean (intercept) | 2.8 | 0.8 | 1.6; 4.0 |
| Background | -0.04 | 0.06 | -0.2; 0.07 |
| Blue-dim | 0.07 | 0.06 | -0.04; 0.2 |
| Yellow-bright | -0.03 | 0.06 | -0.1; 0.08 |

Abbreviations: SD = Standard deviation of the estimate; CI = Credible Interval. For each condition, we report differences from the intercept.

#### PVT: Median Reaction Time (RT; S5)

Analyses revealed extreme evidence in favour of a condition difference during the light exposure with the data being approx. 153 times more likely under the H1 than the H0 (BF_10_ = 153.33). More precisely, there was conclusive evidence in favour of faster median reaction times in the background (mean_background_ = 384.4 ± 35.5 ms) than the yellow-bright (mean_yellow-bright_ = 395.5 ± 32.6 ms; BF_10_ = 29.06) or the blue-dim condition (mean_blue-dim_ = 399.8 ± 39.1 ms; BF_10_ = 260.1). There was only anecdotal (inconclusive) evidence against a difference between the yellow-bright and the blue-dim conditions (BF_10_ = 0.39). Table **17** provides an overview of the condition mean (intercept) and deviations from the intercept sampled from the posterior distribution. There was anecdotal evidence against a condition × time interaction with the data being approx. 2 times more likely under the H0 compared to H1 (BF_10_ = 0.47). For a graphical illustration, see Figure **3-c** and Suppl. Fig. **3**. Suppl. Fig. **4** additionally provides an illustration of the PVT results on evenings 1 and 2 (note this was not included in the analysis plan, wherefore we refrain from statistical analyses).

**Table 17:** Median reaction times during the light exposure in the three light conditions sampled from the posterior distribution.

| Effect | Estimate | SD | 95% CI<br>(lower; upper) |
| --- | --- | --- | --- |
| Condition mean (intercept) | 393.0 | 25.4 | 347.4; 438.3 |
| Background | -8.0 | 2.1 | -12.2; -3.8 |
| Blue-dim | 6.0 | 2.1 | 2.0; 10.1 |
| Yellow-bright | 2.0 | 2.0 | -2.0; 6.0 |

Abbreviations: SD = Standard deviation of the estimate; CI = Credible Interval. For each condition, we report differences from the intercept.

#### PVT: Fastest 10% RTs (S6)

For the 10% fastest reaction times during the light exposure, there was only anecdotal and thus inconclusive evidence in favour of a condition difference with the data being approx. 1.8 times more likely under H1 compared to H0 (BF_10_ = 1.77). Table **18** provides an overview of the condition mean (intercept) and deviations from the intercept sampled from the posterior distribution. Regarding the condition × time interaction, there was anecdotal evidence against H1 (BF_10_ = 0.42). Suppl. Fig. **3** provides an illustration of the results. Suppl. Fig. **4** additionally provides an illustration of the PVT results on evenings 1 and 2 (note this was not included in the analysis plan, wherefore we refrain from statistical analyses).

**Table 18:** Fastest 10% reaction times during the light exposure in the three light conditions sampled from the posterior distribution.

| Effect | Estimate | SD | 95% CI<br>(lower; upper) |
| --- | --- | --- | --- |
| Condition mean (intercept) | 334.1 | 25.5 | 298.0; 367.0 |
| Background | -3.6 | 1.5 | -6.6; -0.7 |
| Blue-dim | 2.8 | 1.5 | -0.01; 5.8 |
| Yellow-bright | 0.7 | 1.4 | -2.0; 3.5 |

Abbreviations: SD = Standard deviation of the estimate; CI = Credible Interval. For each condition, we report differences from the intercept.

#### PVT: Slowest 10% RTs (S7)

For the slowest 10% reaction times during the light exposure, there was very strong evidence in favour of H1, i.e., a condition difference (BF_10_ = 60.9). As for the median reaction times, there was conclusive evidence in favour of reaction times being faster in the background (mean_background_ = 481.6 ± 67.8 ms) compared to the yellow-bright (mean_yellow-bright_ = 511.0 ± 81.6 ms; BF_10_ = 22.6) and the blue-dim conditions (mean_blue-dim_ = 516.9 ± 87.4 ms; BF_10_ = 46.15). However, there was moderate (inconclusive) evidence against a condition difference between the blue-dim and the yellow-bright conditions (BF_10_ = 0.26). Table **19** provides an overview of the condition mean (intercept) and deviations from the intercept sampled from the posterior distribution. There was moderate to strong evidence against a condition × time interaction (BF_10_ = 0.09). For a graphical illustration, see Suppl. Fig. **3**. Suppl. Fig. **4** additionally provides an illustration of the PVT results on evenings 1 and 2 (note this was not included in the analysis plan, wherefore we refrain from statistical analyses).

**Table 19:**
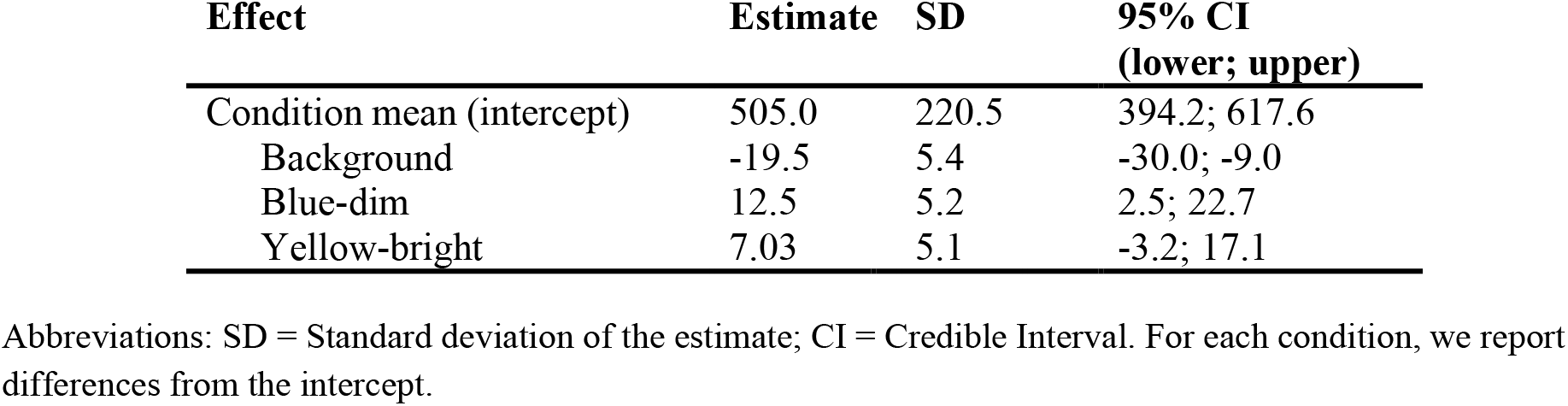
Slowest 10% reaction times during the light exposure in the three light conditions sampled from the posterior distribution.

| Effect | Estimate | SD | 95% CI<br>(lower; upper) |
| --- | --- | --- | --- |
| Condition mean (intercept) | 505.0 | 220.5 | 394.2; 617.6 |
| Background | -19.5 | 5.4 | -30.0; -9.0 |
| Blue-dim | 12.5 | 5.2 | 2.5; 22.7 |
| Yellow-bright | 7.03 | 5.1 | -3.2; 17.1 |

Abbreviations: SD = Standard deviation of the estimate; CI = Credible Interval. For each condition, we report differences from the intercept.

#### EEG-derived Sleep Onset Latency (SLAT; S8)

Analyses yielded inconclusive evidence against a condition difference regarding the onset latency to 10 minutes of continuous sleep. The data were approx. 3 times more likely given the H0 than the H1 (BF_10_ = 0.31; BF_10 rank-based_ = 0.39). The mean latency to 10 minutes of continuous sleep was 17.0 ± 42.2 minutes in the background, 9.4 ± 13.2 minutes in the yellow-bright, and 7.5 ± 4.3 minutes in the blue-dim condition. An overview of the condition means (intercept) and standard deviations sampled from the posterior distribution is provided in Table **20**. Suppl. Fig. **5** provides an illustration of the results. For a comprehensive overview of the sleep data for each light condition and visit, see Suppl. Table **2**.

**Table 20:**
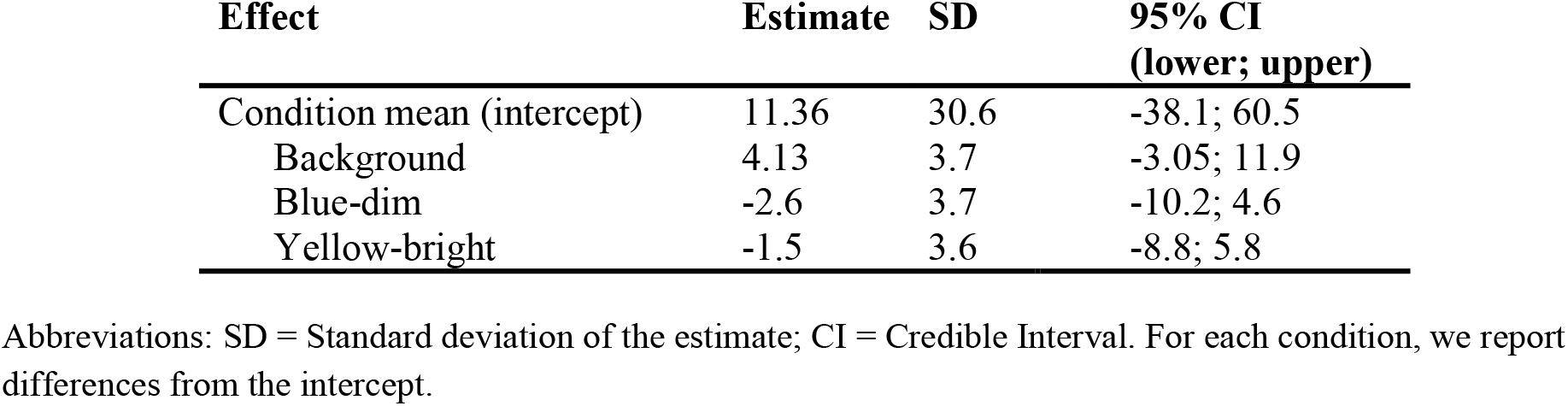
Latency to continuous 10 min of sleep in the three light conditions sampled from the posterior distribution.

#### EEG-derived Slow Wave Activity (SWA; S9)

Note that during two nights, one participant had a very short REM latency (i.e., < 30 min), which would have resulted in a very short NREM part. We thus decided to combine the NREM parts of the detected first and second sleep cycles into one.

Analyses of slow-wave activity (0.5-4.5 Hz) at frontal electrodes F3, Fz, and F4 during the NREM part of the first sleep cycle yielded conclusive evidence against a condition difference (BF_10_ = 0.03; BF_10 rank-based_ = 0.03). Thus, the data was approx. 33 times more likely under the H0 than under H1. Evidence for a time × condition difference remained inconclusive (anecdotal evidence against H0; BF_10_ = 0.42; BF_10 rank-based_ = 0.43). For a graphical illustration, see Figure **3-e**. Table **21** provides an overview of the condition mean (intercept) and deviations from the intercept sampled from the posterior distribution.

**Table 21:** Slow wave activity during the first sleep cycle in the three light conditions sampled from the posterior distribution.

| Effect | Estimate | SD | 95% CI<br>(lower; upper) |
| --- | --- | --- | --- |
| Condition mean (intercept) | 106.0 | 100.4 | -49.7; 260.5 |
| Background | -1.2 | 4.3 | -9.6; 7.1 |
| Blue-dim | -2.0 | 4.3 | -10.4; 6.2 |
| Yellow-bright | 3.3 | 4.3 | -5.2; 11.8 |

Abbreviations: SD = Standard deviation of the estimate; CI = Credible Interval. For each condition, we report differences from the intercept.

### Outcome-Neutral Measurements

#### Melatonin concentrations (ON1)

There was strong evidence against a condition difference in melatonin concentrations before the beginning of the light exposure (BF_10_ = 0.03; BF_10 rank-based_ = 0.03) suggesting that the data was approximately 33 times more likely under the H0 than under the H1. Table **22** provides an overview of the condition mean (intercept) and standard deviations sampled from the posterior distribution. For the condition × time interaction, there was extreme evidence against H1 (BF_10_ = 0.003; BF_10 rank-based_ = 0.003). For a graphical illustration, see Figure **3-b**.

**Table 22:** Melatonin values before the start of the light exposure in the three light conditions sampled from the posterior distribution.

| Effect | Estimate | SD | 95% CI<br>(lower; upper) |
| --- | --- | --- | --- |
| Condition mean (intercept) | 8.2 | 6.6 | -0.9; 17.9 |
| Background | 0.3 | 0.25 | -0.2; 0.8 |
| Blue-dim | -0.1 | 0.25 | -0.6; 0.4 |
| Yellow-bright | -0.1 | 0.25 | -0.6; 0.4 |

Abbreviations: SD = Standard deviation of the estimate; CI = Credible Interval. For each condition, we report differences from the intercept.

#### PVT: Median Reaction Time (RT, ON2)

Before beginning of the light exposure, analyses yielded strong evidence in favour of a condition difference (BF_10_ = 17.42). Follow-up analyses indicated evidence in favour of faster reaction times in the background (mean_background_ = 359.1 ± 36.7 ms) compared to the yellow-bright (mean_yellow-bright_ = 365.2 ± 42.5 ms) or the blue-dim condition (mean_blue-dim_ = 365.8 ± 37.9 ms; BF_10_ = 19.8). There was only moderate inconclusive evidence in favour of a difference between the yellow-bright and the blue-dim conditions (BF_10_ = 5.49). For an overview of the condition mean (intercept) and standard deviations sampled from the posterior distribution please see Table **23**. Regarding a condition × time interaction, there was extreme evidence against H1 with the data being approx. 222 times more likely under the H0 compared to H1 (BF_10_ = 0.004). Figure **3-c** provides a graphical illustration of the results.

**Table 23:**
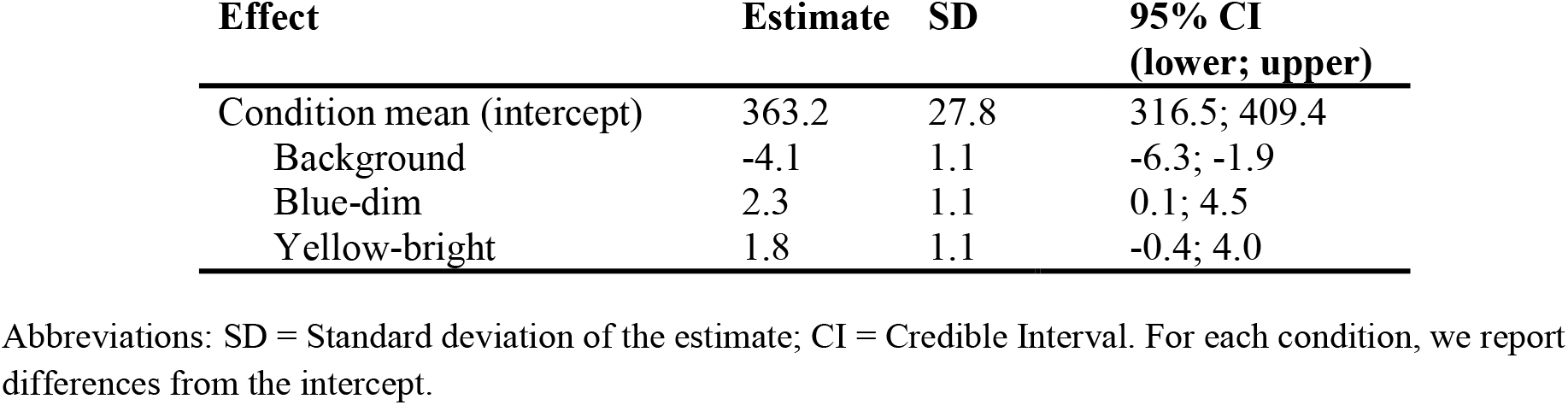
Median reaction times prior to the light exposure in the three light conditions sampled from the posterior distribution.

| Effect | Estimate | SD | 95% CI<br>(lower; upper) |
| --- | --- | --- | --- |
| Condition mean (intercept) | 363.2 | 27.8 | 316.5; 409.4 |
| Background | -4.1 | 1.1 | -6.3; -1.9 |
| Blue-dim | 2.3 | 1.1 | 0.1; 4.5 |
| Yellow-bright | 1.8 | 1.1 | -0.4; 4.0 |

Abbreviations: SD = Standard deviation of the estimate; CI = Credible Interval. For each condition, we report differences from the intercept.

#### Subjective Sleepiness (ON3)

For the subjective sleepiness before the beginning of the light exposure, there was extreme evidence in favour of H1, that is, a condition difference with the data being >100 times more likely under the H1 compared to H0 (BF_10_ = 397569714). More specifically, participants felt more tired in the background (mean_KSS background_ = 6.19 ± 1.6) compared to the yellow-bright condition (mean_KSS yellow-bright_ = 5.56 ± 1.7; BF_10_ = 548934; extreme evidence in favour of H1) and to the blue-dim condition (mean_KSS blue-dim_ = 5.47 ± 1.5; BF_10_ = 321238205; extreme evidence in favour of H1). There was moderate (inconclusive) evidence against a condition difference between the yellow-bright and the blue-dim conditions (BF_10_ = 0.16; moderate evidence against H1). Table **24** provides an overview of the condition means (intercept) for the subjective sleepiness scores and standard deviations sampled from the posterior distribution. Analyses further yielded very strong evidence against a condition × time interaction with the data being approx. 86 times more likely under the H0 than under H1 (BF_10_ = 0.01). For a graphical illustration, see Figure **3-d**.

**Table 24:** Subjective sleepiness scores before the start of the light exposure in the three light conditions sampled from the posterior distribution.

| Effect | Estimate | SD | 95% CI<br>(lower; upper) |
| --- | --- | --- | --- |
| Condition mean (intercept) | 5.7 | 1.7 | 3.23; 8.1 |
| Background | 0.4 | 0.1 | 0.3; 0.6 |
| Blue-dim | -0.3 | 0.1 | -0.4; -0.1 |
| Yellow-bright | -0.2 | 0.1 | -0.3; -0.1 |

Abbreviations: SD = Standard deviation of the estimate; CI = Credible Interval. For each condition, we report differences from the intercept.

#### Brightness (ON4)

There was extreme evidence in favour of a condition difference between the three light exposure conditions prior to the light exposure (BF_10_ = 110.93) with the data being approx. 111 times more likely under the H1 than the H0. The mean perceived brightness (±SD) in the background condition (range: 1-5) was 3.3 ± 0.56, in the blue-dim condition 3.4 ± 0.53, and in the yellow-bright condition 3.5 ± 0.55. More specifically, there was extreme evidence in favour of a condition difference between the background and the yellow-bright conditions (BF_10_ = 752.3), moderate (inconclusive) evidence against a difference between background and the blue-dim conditions (BF_10_ = 0.31) and moderate (inconclusive) evidence in favour of a difference between the blue-dim and yellow-bright conditions (BF_10_ = 7.72). For an overview of the condition mean (intercept) and standard deviations sampled from the posterior distribution please see Table **25**. For the condition × time interaction, there was anecdotal (inconclusive) evidence against an effect (BF_10_ = 0.36).

**Table 25:** Perceived brightness before the start of the light exposure in the three conditions sampled from the posterior distribution.

| Effect | Estimate | SD | 95% CI<br>(lower; upper) |
| --- | --- | --- | --- |
| Condition mean (intercept) | 3.4 | 0.9 | 2.5; 4.3 |
| Background | -0.07 | 0.02 | -0.1; -0.03 |
| Blue-dim | -0.02 | 0.02 | -0.1; 0.02 |
| Yellow-bright | 0.09 | 0.02 | 0.04; 0.1 |

Abbreviations: SD = Standard deviation of the estimate; CI = Credible Interval. For each condition, we report differences from the intercept.

## Discussion

In this Registered Report, we found no conclusive evidence for an effect of calibrated silent-substitution changes in light colour along the blue-yellow axis on the human circadian clock or sleep. In a targeted test of our primary hypothesis, there was no conclusive evidence for differential phase-delaying effects of a 1-h nocturnal light exposure (starting 30 min after habitual bedtime) to constant background/control light, blue-dim, and yellow-bright flickering stimuli using typical evening light levels and constant melanopic excitation across light conditions. Additionally, there were no conclusive differences in melatonin suppression. Thus, we conclude that, even if there is an effect we have missed, the contribution of a postreceptoral channel, where S-cone signals are pitted against a joint L+M signal (i.e., luminance; S-[L+M]), is probably not physiologically relevant to the circadian timing system in healthy humans at night under typical light exposure conditions. Rather, this underscores a likely primary role of melanopsin-containing ipRGCs in mediating these effects as has previously been demonstrated repeatedly ^e.g., 68–71^. In line with this, a recent meta-analysis^72^ concluded that melanopic EDI was the best single predictor for melatonin suppression at light levels above 21 photopic lux. Furthermore, there was no conclusive evidence for differential effects of the calibrated changes in light colour on visual comfort, subjective and objective sleepiness, psychomotor vigilance, EEG-derived sleep onset latency, or homeostatic sleep pressure as assessed by slow-wave activity during the first sleep cycle.

In more detail, the 1-h light exposure starting 30 min after habitual bedtime induced a phase delay in all three light exposure conditions suggesting that the light exposure generally had an effect on the circadian timing system (BFs from post hoc t-tests assessing difference from zero: BF_BKG_ = 59.5; BF_B_ = 14.8; BF_Y_ = 86.8). Given that the unmasked endogenous period length under dim light conditions has been suggested to be about 24.3 hours^73^, it seems unlikely that the observed phase delays can solely be attributed to the lack of morning light. However, evidence for the hypothesis that yellow-bright flickering light would induce a larger phase delay than blue-dim light as well as that flickering light would generally lead to larger shifts than the background condition, remained inconclusive. Our results contrast earlier findings in mice by Mouland and colleagues^74^ who reported that a light stimulus biased towards S-opsin activation thus appearing blue resulted in weaker circadian responses than “yellow” stimuli with a bias towards L-cone activation when rod- and melanopsin activation were held constant. Specifically, in their study, the period length as indicated by the length of the mice’s rest-activity cycles was prolonged under yellow light suggesting a constant stronger phase shift than in the blue condition. Apart from the difference in the studied species, one explanation for the deviating findings could be differences in the duration and timing of the light exposure. In Mouland et al.’s study, mice were exposed to the yellow or blue light at all times during the light phases^74^, whereas we only used a 1-h light exposure at a specific circadian phase (i.e., starting 30 min after habitual bed time). However, earlier findings by Gooley and colleagues^75^ had suggested that cones substantially contribute to effects especially at the beginning of a light exposure and at rather low irradiance levels. Here, participants had been exposed to a narrow-bandwidth 555 nm green light or blue light at 460 nm (10–14 nm half-peak bandwidth) specifically targeting melanopsin for 6.5 hours starting near the onset of nocturnal melatonin secretion (16 irradiance levels covering a broad range of photon densities randomised across participants; 2.52 × 10^11^ − 1.53 × 10^14^ photons/cm^2^/sec). Regarding the phase-resetting response, the authors concluded that the effects of the exposure to green relative to blue light were too large to solely be attributed to melanopsin excitation, suggesting an involvement of the cones. Likewise, both lights were equally effective in suppressing melatonin at the beginning of the exposure, with the importance of melanopsin increasing exponentially with the length of exposure. This notion has recently received further support from a publication resulting from the same study, where the phase-shifting and melatonin suppressing responses were investigated in now six wavelength conditions (420 nm, 460 nm, 480 nm, 507 nm, 555 nm, and 620 nm; photon densities ranging from 2.52 × 10^11^ to 1.53 × 10^14^ photons/cm^2^/sec)^76^. Here, ipRGCs contributed 33%, S-cones 51%, and L+M cones 16% to the phase-shifting effects of the 6.5-h light exposure. Regarding melatonin suppression, during the first quarter of the 6.5-h light exposure (i.e., 97.5 min), S-cones and L+M cones substantially contributed to melatonin suppression (51% and 47% contributions of the respective spectral sensitivity curves to the overall sensitivity) while ipRGCs only contributed 2%. As in Gooley et al. (2010)^75^, the contribution of ipRGCs and thus melanopsin significantly increased across time. Thus, we argue that a 1-h exposure specifically targeting cones should have been sufficient to produce differential effects in the present study. Furthermore, several investigations of the interaction between circadian phase and the magnitude and direction of the effects of light exposure have suggested increased sensitivity of the circadian timing system at the phase of exposure to bright^77, 78^ and blue light^79^. Nevertheless, future studies will need to evaluate the effects of calibrated colour changes along the blue-yellow axis at different circadian phases, and different mean light levels, and to investigate whether there is a specific sensitivity to calibrated changes along the blue-yellow dimension, for instance specifically during the twilight hours.

Conclusive condition differences were limited to differences in psychomotor vigilance, that is, reaction times on the Psychomotor Vigilance Task (PVT) and subjective sleepiness as assessed with the Karolinska Sleepiness Scale (KSS). Considering median reaction times, participants were faster in the background condition than in the blue-dim or the yellow-bright conditions before as well as during the light exposure. During light exposure, this effect was also visible when looking at the 10% slowest responses. Especially because reaction time differences were not limited to the 1-h light exposure but present even before, this most likely seems an effect of increased motivation during the first experimental visit (always “background” condition). Interestingly and somewhat contradictory, the faster reaction times in the background condition coincided with higher subjective sleepiness ratings. This difference was however limited to the time before the light exposure. We speculate that may have resulted from participants not knowing what to expect and how they would react to the prolonged wake time and the laboratory protocol, which may have been perceived as tiring.

Limitations of the study include the fact that the order of the conditions were not completely randomised, but the background condition was always the first one. This had logistical reasons as randomising three conditions would have resulted in six potential condition orders. Given the results from the Bayes Factor Design Analysis^36^, such an approach would possibly have required us to obtain data from 2 × 12 participants (i.e., 2 × 6 men and 6 women), which would have exceeded the financial and resource limit of this project. Besides this, the fact that the light exposure took place between 30 min and 1 h 30 min after habitual bedtime may have resulted in homeostatic sleep pressure overriding more subtle effects of the different light exposure conditions. However, other studies investigating the effects of light, where participants likewise went to sleep well after habitual bedtime had still found differences between monochromatic light exposure conditions on for instance slow wave activity at the beginning of the night^25, 80^, self-reported sleepiness^81^, and psychomotor vigilance^81^. Furthermore, we did not objectively assess light history prior to the arrival at the lab, which is known to affect the response to a subsequent light stimulus^82, 83^. However, evidence for differences between the conditions in self-reported time spent under the open sky prior to arriving at the lab (BF_10_ = 0.33; BF_10 rank-based_ = 0.16), or self-reported lighting conditions outside on that day during the time they had spent under the open sky remained inconclusive (BF_10_ = 0.16; 10-point Likert scale from “very dull day” to “bright summer’s day”). Additionally, participants’ visits were scheduled within 3 weeks and took place on the same day of the week contributing to a maximal stability of ambient lighting conditions as well as individual schedules and thus light history. Furthermore, we did not control pupil size, which may have led to small changes in retinal irradiance, but we consider it implausible that this could explain the lack of a conclusive effect. Lastly, we assumed that rods do not contribute to circadian effects under photopic conditions. Thus, rhodopic effects were not constant across conditions (Supplemental Material **S3** and **S4**). Data obtained in mice lacking functional cones or mice in which rods were the only photoreceptors suggest that rods support circadian photoentrainment even at higher light levels (i.e., 500 lux)^84^, although the effects on the SCN may be rather small^85^. Recent data suggests, that the same holds true for humans living with functional achromatopsia, i.e., individuals lacking functional cones^86^. However, the relative contributions of rod and melanopsin contributions are unknown and some degree of adaptation of the circadian system in this congenital condition is likely^87, 88^. Future studies should carefully disentangle the unique contributions of the rods to circadian photoreception across light levels.

In summary, we found no conclusive evidence for an effect of calibrated changes in light colour along the blue-yellow axis at constant melanopic excitation on the human circadian system, psychomotor vigilance, sleepiness, or sleep (i.e., latency to 10 min of continuous sleep, slow wave activity during the first sleep cycle) for a 1-h light exposure starting 30 min after habitual bedtime. This seems to underscore the primary role of melanopsin-containing ipRGCs in mediating these effects that have repeatedly been reported in the literature. From a more practical perspective, it seems that the human circadian clock is relatively insensitive to shifts in light colour towards warmer colour temperatures at constant melanopic excitation. Smartphones and other displays with night-shift modes typically reduce colour *and* melanopic excitation in a yoked fashion, and our study provides evidence that any effects seen in night-shift mode may be due to the reduction in melanopic excitation. As a large body of literature convincingly suggests that short-wavelength proportions of light should be reduced in the evening^37^ to prevent decreases in sleepiness and a phase-delay^37, 68, 89, 90^, we therefore encourage users of devices with background-lit displays (i.e., smartphones, tablets, computer screens) to make use of built-in software or apps such as f.lux^91^ in the evenings and during the night. In the future, tech companies may also opt to use metameric light that allows to reduce short-wavelength proportions without a change in perceived colour. Recently, Schöllhorn and colleagues^69^ showed that low-melanopic light can indeed mitigate the unwanted effects of screen use at night, confirming the primary role of melanopsin photoreception in setting our circadian system by light.

## Data availability

All data generated in this study (including laboratory logs) is available in anonymised and deidentified form on FigShare (Data: https://doi.org/10.6084/m9.figshare.23578698; Laboratory Log: https://doi.org/10.6084/m9.figshare.23578695).

## Code availability

Code to simulate and analyse data and implement statistical models is available on a dedicated GitHub repository (https://github.com/ChristineBlume/Effects-of-calibrated-blue-yellow-changes-in-light-on-the-human-circadian-clock/) which will be archived as a snapshot on FigShare. The simulation code was shared with reviewers via the journal in the Stage 1 submission and was be made public at Stage 2.

## Protocol registration

The approved Stage 1 protocol can be found here (https://springernature.figshare.com/articles/journal_contribution/Effects_of_calibrated_blue-yellow_S_L_M_S_L_M_changes_in_light_on_the_human_circadian_clock_Registered_Report_Stage_1_Protocol_/13050215), a table of the laboratory protocol can be found here (https://figshare.com/articles/dataset/Protocol/23578704).

## Ethical approval

Approval for this study was granted from the Ethikkommission Nordwest- und Zentralschweiz (EKNZ) with approval number 2020-02037. The study was conducted in accordance with Swiss law and the Declaration of Helsinki. It was registered as a clinical trial (DRKS00023603).

## Author Contributions

CB and MS conceptualised the project and wrote the Stage 1 Registered Report supported by CC. CB and MS acquired the funding. CB acquired the data, conducted the analyses, and wrote the first draft of the full manuscript supported by MS. IS and HCS greatly supported the data acquisition. All authors provided a critical review of the manuscript and approved the submitted version.

## Funding

CB was supported by a scholarship from the University of Basel’s research fund for excellent early career researchers as well as an Ambizione grant from the Swiss National Science Foundation awarded to her. Additionally, the study was funded by a grant from the Forschungsförderungsfonds of the Psychiatric Hospital of the University of Basel (UPK) awarded to MS. During parts of this work, MS was supported by a Sir Henry Wellcome Postdoctoral Fellowship (Wellcome Trust, 204686/Z/16/Z) and Linacre College, University of Oxford (Biomedical Sciences Junior Research Fellowship).

## Competing interest

The authors have no competing interests related to this publication. For full transparency, the authors declare the following non-competing interests. Related to lighting, MS is currently an unpaid member of CIE Technical Committee TC 1-98 (“A Roadmap Toward Basing CIE Colorimetry on Cone Fundamentals”). MS was an unpaid advisor to the Division Reportership DR 6-45 of Division 3 (“Publication and maintenance of the CIE S026 Toolbox”) and a member of the CIE Joint Technical Committee 9 on the definition of CIE S 026:2018. Since 2020, MS is an elected Member of the Daylight Academy, an unpaid member of the Board of Advisors of the Center for Environmental Therapeutics. CC has had the following commercial interests related to lighting: honoraria, travel, accommodation and/or meals for invited keynote lectures, conference presentations or teaching from Toshiba Materials, Velux, Firalux, Lighting Europe, Electrosuisse, Novartis, Roche, Elite, Servier, and WIR Bank. CC is an elected member of the Daylight Academy. CB has had the following commercial interests related to sleep and/or light: honoraria for invited talks and workshops from IKEA, F.A. Hoffmann-La Roche AG, L’Oréal, and Vattenfall. CB is an elected member of the Daylight Academy.

## Acknowledgments

First, we thank all participants, who volunteered to participate in the project. The data acquisition has mainly been done by an amazing team of helping hands, mainly interns, and master students. We are incredibly grateful to Zarah Butt, who was also the study coordinator, Saira Bogazlyianlioglu, Lina Fricke, Alexandra von Gatterburg, Yvonne Hao, Bernarda Knezevic, Ann-Sophie Loock, Ilenia Messina, Joé Miller, Maria Vettiger, Leticia Vilela, Patrick Weiss, Kerstin Wieczorek, Ronja Zeugin, and Tamara Zumbrunn. Thank you for your team spirit, your flexibility and for making this project possible. We also thank Fatemeh Fazlali for her help with the calibration of the display, Mirjam Münch for sharing her knowledge on the determination of DLMOs using the Hockeystick algorithm and Jakob Weber from NovoLytiX for constant support with the melatonin assays. Last, we also thank the study physicians Martin P. Meyer, and Christian Epple as well as Marco Cattaneo, statistician at the Department of Clinical Research of the University of Basel, for their valuable support and advice.

## Supplementary material

**Supplementary Material S1.** Example protocol for the experimental assessments (assumed habitual bedtime 00:00).

**Supplementary Material S2.** Suppl. figures, methods, results, tables.

**Supplementary Material S3.** Table with irradiance-derived measures.

**Supplementary Material S4.** Table with radiance-derived measures.

**Supplementary Material S5.** Spectral distribution of irradiance for the different settings and light exposure conditions.

**Supplementary Material S6.** Spectral distribution of radiance for the three light exposure conditions.

---

**Picture 1.**
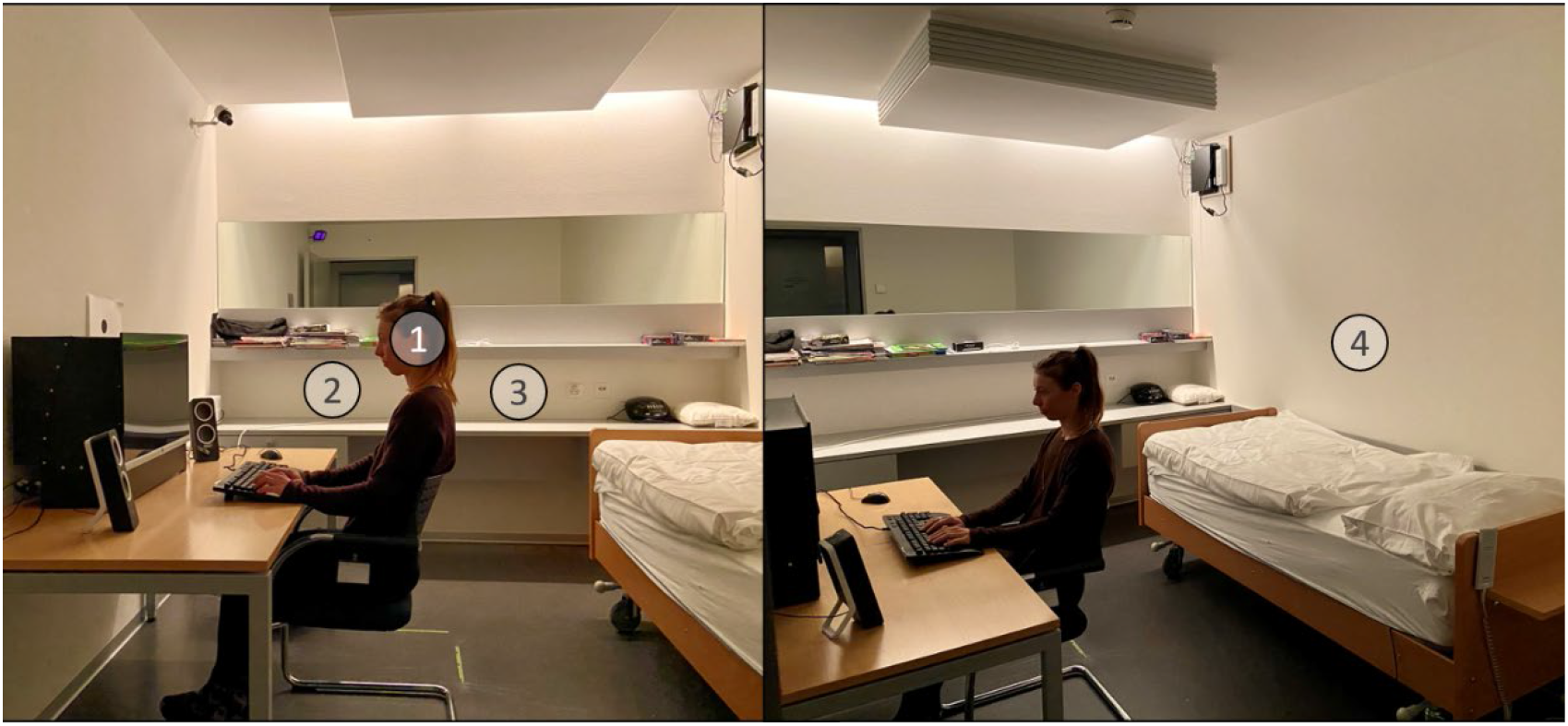
Laboratory setup. Photo of the laboratory setup and illustrations of the four locations, where measurements of ambient illuminance were taken from the observer’s point of view. The numbers in the picture correspond to (1) “at desk with display”, (2) “on the sofa facing away from the ambient light”, (3) “at the desk facing the surface of the desk”, and (4) “on the bed, facing the opposite wall”. Participants spent most of the time during the day and in the evenings at locations 1 and 2.

## Notes

### Competing Interest Statement

The authors have declared no competing interest.

## References

1 Blume, C., Garbazza, C. & Spitschan, M. Effects of light on human circadian rhythms, sleep and mood. Somnologie (Berl) 23, 147–156 (2019). https://doi.org:10.1007/s11818-019-00215-x

2 Thapan, K., Arendt, J. & Skene, D. J. An action spectrum for melatonin suppression: evidence for a novel non-rod, non-cone photoreceptor system in humans. J Physiol 535, 261–267 (2001). https://doi.org:10.1111/j.1469-7793.2001.t01-1-00261.x

3 Brainard, G. C. et al. Action spectrum for melatonin regulation in humans: evidence for a novel circadian photoreceptor. J Neurosci 21, 6405–6412 (2001).

4 Prayag, A. S., Najjar, R. P. & Gronfier, C. Melatonin suppression is exquisitely sensitive to light and primarily driven by melanopsin in humans. J Pineal Res 66, e12562 (2019). https://doi.org:10.1111/jpi.12562

5 Wright, H. R. & Lack, L. C. Effect of light wavelength on suppression and phase delay of the melatonin rhythm. Chronobiol Int 18, 801–808 (2001). https://doi.org:10.1081/cbi-100107515

6 Gooley, J. J. et al. Spectral responses of the human circadian system depend on the irradiance and duration of exposure to light. Sci Transl Med 2, 31ra33 (2010). https://doi.org:10.1126/scitranslmed.3000741

7 Spitschan, M. Melanopsin contributions to non-visual and visual function. Curr Opin Behav Sci 30, 67–72 (2019). https://doi.org:10.1016/j.cobeha.2019.06.004

8 Dacey, D. M. et al. Melanopsin-expressing ganglion cells in primate retina signal colour and irradiance and project to the LGN. Nature 433, 749–754 (2005). https://doi.org:10.1038/nature03387

9 Patterson, S. S., Kuchenbecker, J. A., Anderson, J. R., Neitz, M. & Neitz, J. A Color Vision Circuit for Non-Image-Forming Vision in the Primate Retina. Current Biology (2020). https://doi.org:10.1016/j.cub.2020.01.040

10 Spitschan, M., Jain, S., Brainard, D. H. & Aguirre, G. K. Opponent melanopsin and S-cone signals in the human pupillary light response. Proc Natl Acad Sci U S A 111, 15568–15572 (2014). https://doi.org:10.1073/pnas.1400942111

11 Woelders, T. et al. Melanopsin- and L-cone-induced pupil constriction is inhibited by S- and M-cones in humans. Proc Natl Acad Sci U S A 115, 792–797 (2018). https://doi.org:10.1073/pnas.1716281115

12 Spitschan, M., Lazar, R., Yetik, E. & Cajochen, C. No evidence for an S cone contribution to acute neuroendocrine and alerting responses to light. Curr Biol 29, R1297–R1298 (2019). https://doi.org:10.1016/j.cub.2019.11.031

13 Czeisler, C. A. et al. Suppression of melatonin secretion in some blind patients by exposure to bright light. N Engl J Med 332, 6–11 (1995). https://doi.org:10.1056/NEJM199501053320102

14 Hull, J. T., Czeisler, C. A. & Lockley, S. W. Suppression of Melatonin Secretion in Totally Visually Blind People by Ocular Exposure to White Light: Clinical Characteristics. Ophthalmology 125, 1160–1171 (2018). https://doi.org:10.1016/j.ophtha.2018.01.036

15 Jacobs, G. H. The discovery of spectral opponency in visual systems and its impact on understanding the neurobiology of color vision. J Hist Neurosci 23, 287–314 (2014). https://doi.org:10.1080/0964704X.2014.896662

16 Kaiser, P. K. & Boynton, R. M. Human color vision. 2nd edn, (Optical Society of America, 1996).

17 Stockman, A. & Brainard, D. H. (ed M Bass) 11.11–11.104 (McGraw-Hill, 2010).

18 Krauskopf, J., Williams, D. R. & Heeley, D. W. Cardinal directions of color space. Vision Res 22, 1123–1131 (1982). https://doi.org:10.1016/0042-6989(82)90077-3

19 Figueiro, M. G., Bullough, J. D., Parsons, R. H. & Rea, M. S. Preliminary evidence for spectral opponency in the suppression of melatonin by light in humans. Neuroreport 15, 313–316 (2004). https://doi.org:10.1097/00001756-200402090-00020

20 Spitschan, M., Aguirre, G. K., Brainard, D. H. & Sweeney, A. M. Variation of outdoor illumination as a function of solar elevation and light pollution. Sci Rep 6, 26756 (2016). https://doi.org:10.1038/srep26756

21 Woelders, T., Wams, E. J., Gordijn, M. C. M., Beersma, D. G. M. & Hut, R. A. Integration of color and intensity increases time signal stability for the human circadian system when sunlight is obscured by clouds. Sci Rep 8, 15214 (2018). https://doi.org:10.1038/s41598-018-33606-5

22 Spitschan, M., Lucas, R. J. & Brown, T. M. Chromatic clocks: Color opponency in non-image-forming visual function. Neurosci Biobehav Rev 78, 24–33 (2017). https://doi.org:10.1016/j.neubiorev.2017.04.016

23 Roenneberg, T. & Foster, R. G. Twilight times: light and the circadian system. Photochem Photobiol 66, 549–561 (1997). https://doi.org:10.1111/j.1751-1097.1997.tb03188.x

24 Mouland, J. W., Martial, F., Watson, A., Lucas, R. J. & Brown, T. M. Cones Support Alignment to an Inconsistent World by Suppressing Mouse Circadian Responses to the Blue Colors Associated with Twilight. Curr Biol 29, 4260–4267 e4264 (2019). https://doi.org:10.1016/j.cub.2019.10.028

25 Chellappa, S. L. et al. Acute exposure to evening blue-enriched light impacts on human sleep. Journal of sleep research 22, 573–580 (2013).

26 Chellappa, S. L. et al. Non-visual effects of light on melatonin, alertness and cognitive performance: can blue-enriched light keep us alert? PloS one 6, e16429 (2011).

27 Webster, M. A. & Wilson, J. A. Interactions between chromatic adaptation and contrast adaptation in color appearance. Vision Research 40, 3801–3816 (2000). https://doi.org:10.1016/s0042-6989(00)00238-8

28 Walraven, J., Enroth-Cugell, C., Hood, D. C., MacLeod, D. I. A. & Schnapf, J. L. in Visual perception: The neurophysiological foundations (eds L. Spillman & J. S. Werner) (Academic Press, 1990).

29 von Kries, J. in Sources of color science (ed D. L. MacAdam) (MIT Pres, 1970).

30 Rinner, O. & Gegenfurtner, K. R. Time course of chromatic adaptation for color appearance and discrimination. Vision Research 40, 1813–1826 (2000). https://doi.org:10.1016/s0042-6989(00)00050-x

31 Jameson, D. & Hurvich, L. M. in Visual Psychophysics. Handbook of Sensory Physiology*, vol 7 / 4* (eds D. Jameson & L.M. Hurvich) 568–581 (Springer, 1972).

32 Webster, M. A. & Tregillus, K. E. M. Visualizing Visual Adaptation. J Vis Exp (2017). https://doi.org:10.3791/54038

33 Hood, D. C. & Finkelstein, M. A. in Visual Psychophysics: Its Physiological Basis (eds K. R. Boff, L. Kaufman, & J. P. Thomas) (Academic Press, 1986).

34 Najjar, R. P. & Zeitzer, J. M. Temporal integration of light flashes by the human circadian system. J Clin Invest 126, 938–947 (2016). https://doi.org:10.1172/JCI82306

35 Schonbrodt, F. D., Wagenmakers, E. J., Zehetleitner, M. & Perugini, M. Sequential hypothesis testing with Bayes factors: Efficiently testing mean differences. Psychol Methods 22, 322–339 (2017). https://doi.org:10.1037/met0000061

36 BFDA: An R package for Bayes factor design analysis v. 0.3 (2018).

37 Chang, A.-M., Aeschbach, D., Duffy, J. F. & Czeisler, C. A. Evening use of light-emitting eReaders negatively affects sleep, circadian timing, and next-morning alertness. Proceedings of the National Academy of Sciences 112, 1232–1237 (2015).

38 Cote, K. A. et al. CNS arousal and neurobehavioral performance in a short-term sleep restriction paradigm. J Sleep Res 18, 291–303 (2009). https://doi.org:10.1111/j.1365-2869.2008.00733.x

39 Basner, M. & Dinges, D. F. Maximizing sensitivity of the psychomotor vigilance test (PVT) to sleep loss. Sleep 34, 581–591 (2011).

40 Dinges, D. F. & Powell, J. W. Microcomputer analyses of performance on a portable, simple visual RT task during sustained operations. Behavior Research Methods, Instruments, & Computers 17, 652–655 (1985). https://doi.org:10.3758/bf03200977

41 Dorrian, J., Rogers, N. & Dinges, D. (New York, Marcel Dekker, 2005).

42 Münch, M. et al. Effects on subjective and objective alertness and sleep in response to evening light exposure in older subjects. Behavioural brain research 224, 272–278 (2011).

43 Xu, J., Pokorny, J. & Smith, V. C. Optical density of the human lens. J Opt Soc Am A Opt Image Sci Vis 14, 953–960 (1997). https://doi.org:10.1364/josaa.14.000953

44 Pokorny, J., Smith, V. C. & Lutze, M. Aging of the human lens. Appl Opt 26, 1437–1440 (1987). https://doi.org:10.1364/AO.26.001437

45 Roenneberg, T., Wirz-Justice, A. & Merrow, M. Life between Clocks: Daily Temporal Patterns of Human Chronotypes. Journal of Biological Rhythms 18, 80–90 (2003). https://doi.org:10.1177/0748730402239679

46 Mollon, J. & Reffin, J. A computer-controlled colour vision test that combines the principles of Chibret and Stilling. J Physiol 414 (1989).

47 Saletu, B., Wessely, P., Grünberger, J. & Schultes, M. Erste klinische Erfahrungen mit einem neuen schlafanstoßenden Benzodiazepin, Cinolazepam, mittels eines Selbstbeurteilungsbogens für Schlaf-und Aufwachqualität (SSA). Neuropsychiatrie 1, 169–176 (1987).

48 Åkerstedt, T. & Gillberg, M. Subjective and objective sleepiness in the active individual. International Journal of Neuroscience 52, 29–37 (1990).

49 Mifflin, M. D. et al. A new predictive equation for resting energy expenditure in healthy individuals. Am J Clin Nutr 51, 241–247 (1990). https://doi.org:10.1093/ajcn/51.2.241

50 American Academy of Sleep Medicine & Iber, C. The AASM manual for the scoring of sleep and associated events: rules, terminology and technical specifications. (American Academy of Sleep Medicine, 2007).

51 Lasauskaite, R. & Cajochen, C. Influence of lighting color temperature on effort-related cardiac response. Biol Psychol 132, 64–70 (2018). https://doi.org:10.1016/j.biopsycho.2017.11.005

52 Lasauskaite, R., Hazelhoff, E. M. & Cajochen, C. Four minutes might not be enough for light colour temperature to affect sleepiness, mental effort, and light ratings. Lighting Research & Technology 51, 1128–1138 (2018). https://doi.org:10.1177/1477153518796700

53 Spitschan, M. et al. How to Report Light Exposure in Human Chronobiology and Sleep Research Experiments. Clocks Sleep 1, 280–289 (2019). https://doi.org:10.3390/clockssleep1030024

54 R: A Language and Environment for Statistical Computing (R Foundation for Statistical Computing, Vienna, Austria, 2015).

55 Oostenveld, R., Fries, P., Maris, E. & Schoffelen, J.-M. FieldTrip: open source software for advanced analysis of MEG, EEG, and invasive electrophysiological data. Computational Intelligence and Neuroscience 2011 (2010).

56 Danilenko, K. V., Verevkin, E. G., Antyufeev, V. S., Wirz-Justice, A. & Cajochen, C. The hockey-stick method to estimate evening dim light melatonin onset (DLMO) in humans. Chronobiology International 31, 349–355 (2014). https://doi.org:10.3109/07420528.2013.855226

57 Anderer, P. et al. An E-health solution for automatic sleep classification according to Rechtschaffen and Kales: validation study of the Somnolyzer 24× 7 utilizing the Siesta database. Neuropsychobiology 51, 115–133 (2005).

58 Anderer, P. et al. Computer-Assisted Sleep Classification according to the Standard of the American Academy of Sleep Medicine: Validation Study of the AASM Version of the Somnolyzer 24 × 7. Neuropsychobiology 62, 250–264 (2010).

59 Feinberg, I. & Floyd, T. Systematic trends across the night in human sleep cycles. Psychophysiology 16, 283–291 (1979).

60 Blume, C. & Cajochen, C. ‘SleepCycles’ package for R - A free software tool for the detection of sleep cycles from sleep staging. MethodsX 8, 101318 (2021). https://doi.org:https://doi.org/10.1016/j.mex.2021.101318

61 Bates, D., Mächler, M., Bolker, B. & Walker, S. Fitting Linear Mixed-Effects Models Using lme4. 2015 67, 48 (2015). https://doi.org:http://dx.doi.org/10.18637/jss.v067.i01

62 Rouder, J. N., Morey, R. D., Speckman, P. L. & Province, J. M. Default Bayes factors for ANOVA designs. Journal of Mathematical Psychology 56, 356–374 (2012). https://doi.org:10.1016/j.jmp.2012.08.001

63 Roach, G. D., Dawson, D. & Lamond, N. Can a Shorter Psychomotor Vigilance Task Be Usedas a Reasonable Substitute for the Ten-Minute Psychomotor Vigilance Task? Chronobiology International 23, 1379–1387 (2006). https://doi.org:10.1080/07420520601067931

64 Basner, M., Mollicone, D. & Dinges, D. F. Validity and sensitivity of a brief psychomotor vigilance test (PVT-B) to total and partial sleep deprivation. Acta Astronautica 69, 949–959 (2011). https://doi.org:https://doi.org/10.1016/j.actaastro.2011.07.015

65 Schollhorn, I. et al. Melanopic irradiance defines the impact of evening display light on sleep latency, melatonin and alertness. Commun Biol 6, 228 (2023). https://doi.org:10.1038/s42003-023-04598-4

66 Proß, A. Development of a method for display lighting supporting the human circadian system. (Fraunhofer Verlag, 2019).

67 Berry, R. B. et al. The AASM manual for the scoring of sleep and associated events. Rules, Terminology and Technical Specifications, Darien, Illinois, American Academy of Sleep Medicine 176, 2012 (2012).

68 Allen, A. E., Hazelhoff, E. M., Martial, F. P., Cajochen, C. & Lucas, R. J. Exploiting metamerism to regulate the impact of a visual display on alertness and melatonin suppression independent of visual appearance. Sleep 41 (2018). https://doi.org:10.1093/sleep/zsy100

69 Schöllhorn, I. et al. Melanopic irradiance defines the impact of evening display light on sleep latency, melatonin and alertness. Communications Biology 6, 228 (2023). https://doi.org:10.1038/s42003-023-04598-4

70 Santhi, N. et al. The spectral composition of evening light and individual differences in the suppression of melatonin and delay of sleep in humans. Journal of pineal research 53, 47–59 (2012).

71 Najjar, R. P., Prayag, A. S. & Gronfier, C. Melatonin suppression by light involves different retinal photoreceptors in young and older adults. bioRxiv, 2023.2006.2002.543372 (2023). https://doi.org:10.1101/2023.06.02.543372

72 Giménez, M. C. et al. Predicting melatonin suppression by light in humans: Unifying photoreceptor-based equivalent daylight illuminances, spectral composition, timing and duration of light exposure. Journal of Pineal Research n/a, e12786 (2022). https://doi.org:https://doi.org/10.1111/jpi.12786

73 Middleton, B., Arendt, J. & Stone, B. M. Complex effects of melatonin on human circadian rhythms in constant dim light. Journal of biological rhythms 12, 467–477 (1997).

74 Mouland, J. W., Martial, F., Watson, A., Lucas, R. J. & Brown, T. M. Cones Support Alignment to an Inconsistent World by Suppressing Mouse Circadian Responses to the Blue Colors Associated with Twilight. Current Biology 29, 4260–4267. e4264 (2019).

75 Gooley, J. J. et al. Spectral responses of the human circadian system depend on the irradiance and duration of exposure to light. Science translational medicine 2, 31ra33–31ra33 (2010). https://doi.org:10.1126/scitranslmed.3000741

76 St Hilaire, M. A., et al. The spectral sensitivity of human circadian phase resetting and melatonin suppression to light changes dynamically with light duration. Proceedings of the National Academy of Sciences 119, e2205301119 (2022). https://doi.org:doi:10.1073/pnas.2205301119

77 St Hilaire, M. A., et al. Human phase response curve to a 1 h pulse of bright white light. The Journal of Physiology 590, 3035–3045 (2012). https://doi.org:10.1113/jphysiol.2012.227892

78 Khalsa, S. B. S., Jewett, M. E., Cajochen, C. & Czeisler, C. A. A Phase Response Curve to Single Bright Light Pulses in Human Subjects. The Journal of Physiology 549, 945–952 (2003). https://doi.org:10.1113/jphysiol.2003.040477

79 Revell, V. L., Molina, T. A. & Eastman, C. I. Human phase response curve to intermittent blue light using a commercially available device. The Journal of Physiology 590, 4859–4868 (2012). https://doi.org:https://doi.org/10.1113/jphysiol.2012.235416

80 Münch, M. et al. Wavelength-dependent effects of evening light exposure on sleep architecture and sleep EEG power density in men. *American Journal of Physiology - Regulatory*, Integrative and Comparative Physiology 290, R1421–R1428 (2006). https://doi.org:10.1152/ajpregu.00478.2005

81 Lockley, S. W. et al. Short-Wavelength Sensitivity for the Direct Effects of Light on Alertness, Vigilance, and the Waking Electroencephalogram in Humans. Sleep 29, 161–168 (2006). https://doi.org:10.1093/sleep/29.2.161

82 Chang, A.-M., Scheer, F. A. J. L., Czeisler, C. A. & Aeschbach, D. Direct Effects of Light on Alertness, Vigilance, and the Waking Electroencephalogram in Humans Depend on Prior Light History. Sleep 36, 1239–1246 (2013). https://doi.org:10.5665/sleep.2894

83 Hébert, M., Martin, S. K., Lee, C. & Eastman, C. I. The effects of prior light history on the suppression of melatonin by light in humans. Journal of Pineal Research 33, 198–203 (2002). https://doi.org:10.1034/j.1600-079X.2002.01885.x

84 Altimus, C. M. et al. Rod photoreceptors drive circadian photoentrainment across a wide range of light intensities. Nature Neuroscience 13, 1107–1112 (2010). https://doi.org:10.1038/nn.2617

85 Lucas, R. J., Lall, G. S., Allen, A. E. & Brown, T. M. in Progress in Brain Research Vol. 199 (eds Andries Kalsbeek, Martha Merrow, Till Roenneberg, & Russell G. Foster) 1-18 (Elsevier, 2012).

86 Spitschan, M., Garbazza, C., Kohl, S. & Cajochen, C. Sleep and circadian phenotype in people without cone-mediated vision. bioRxiv, 2020.2006.2002.129502 (2021). https://doi.org:10.1101/2020.06.02.129502

87 Giménez, M. C., Beersma, D. G. M., Bollen, P., van der Linden, M. L. & Gordijn, M. C. M. Effects of a chronic reduction of short-wavelength light input on melatonin and sleep patterns in humans: Evidence for adaptation. Chronobiology International 31, 690–697 (2014). https://doi.org:10.3109/07420528.2014.893242

88 Najjar, R. P. et al. Aging of Non-Visual Spectral Sensitivity to Light in Humans: Compensatory Mechanisms? PloS one 9, e85837 (2014). https://doi.org:10.1371/journal.pone.0085837

89 Cajochen, C. et al. High sensitivity of human melatonin, alertness, thermoregulation, and heart rate to short wavelength light. The journal of clinical endocrinology & metabolism 90, 1311–1316 (2005). https://doi.org:10.1210/jc.2004-0957

90 Lockley, S. W., Brainard, G. C. & Czeisler, C. A. High Sensitivity of the Human Circadian Melatonin Rhythm to Resetting by Short Wavelength Light. The Journal of Clinical Endocrinology & Metabolism 88, 4502–4505 (2003). https://doi.org:10.1210/jc.2003-030570

91 f.lux (2023).

92 Webster, M. A. & Mollon, J. D. The influence of contrast adaptation on color appearance. Vision Res 34, 1993–2020 (1994). https://doi.org:10.1016/0042-6989(94)90028-0

93 Cao, D., Nicandro, N. & Barrionuevo, P. A. A five-primary photostimulator suitable for studying intrinsically photosensitive retinal ganglion cell functions in humans. J Vis 15, 15 11 27 (2015). https://doi.org:10.1167/15.1.27

94 luox: Platform for calculating quantities related to light and lighting (2021).

