## Supplemental Material for "Effects of calibrated blue–yellow (–S+[L+M], +S–[L+M]) changes in light on the human circadian clock"

#### *Supplemental Material S3.*

Christine Blume[1,2][0000-0003-2328-9612]§, Christian Cajochen[1,2] [0000-0003-2699-7171], Isabel Schöllhorn[1,2][0000-0001-9067-1614], Helen C. Slawik[3], Manuel Spitschan[4,5,6][0000-0002-8572-9268]

[1]Centre for Chronobiology, Psychiatric Hospital of the University of Basel, Basel, Switzerland (institution where the work was performed)

[2]Research Platform Molecular and Cognitive Neurosciences, University of Basel, Basel, Switzerland

[3]Translational Sensory and Circadian Neuroscience, Max Planck Institute for Biological Cybernetics, Tübingen, Germany

[4]TUM Department of Sport and Health Sciences (TUM SG), Technical University of Munich, Munich, Germany

[5]Psychiatric Hospital of the University of Basel, Basel, Switzerland

**Supplemental Figures**

**Secondary Outcomes.**

**Melatonin Concentrations (S1).**

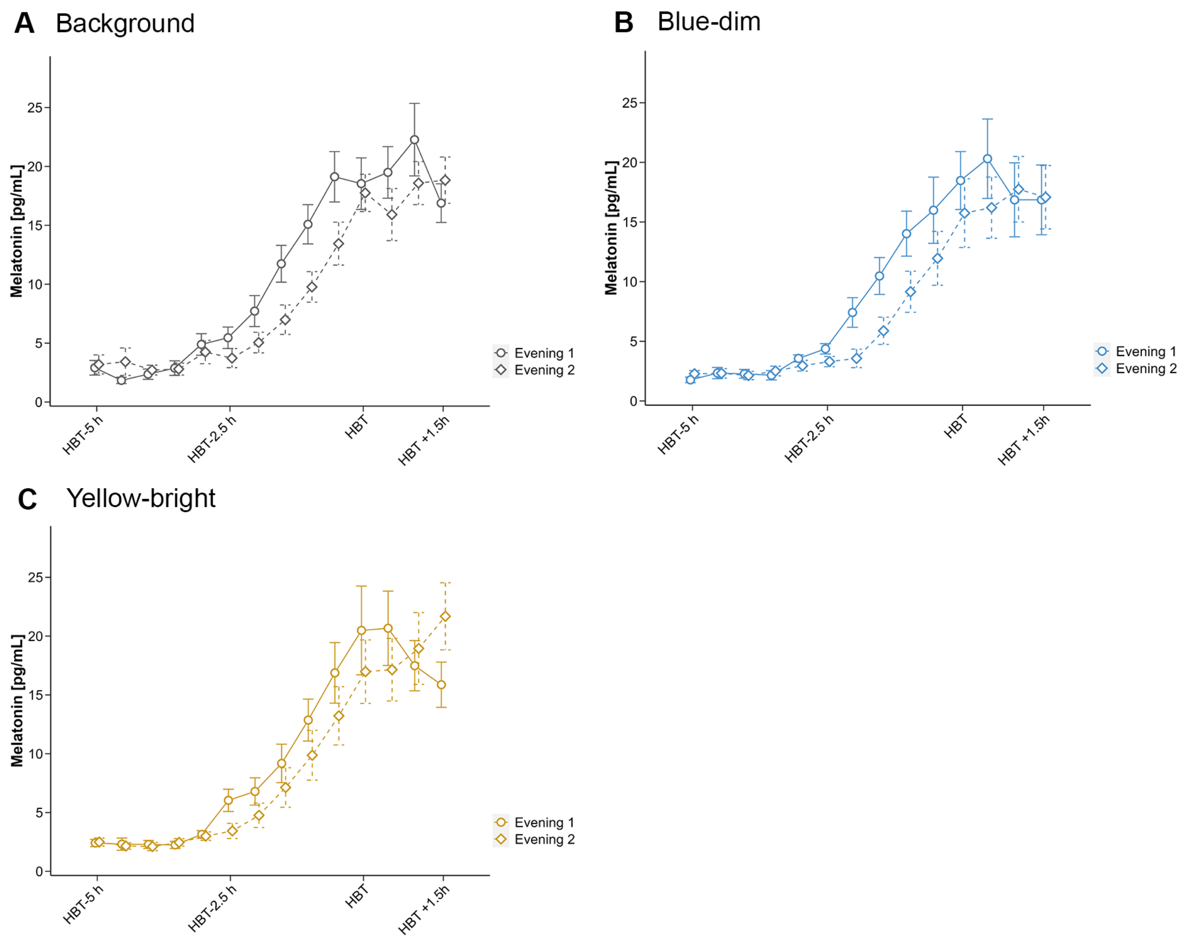


**Supplemental Figure 1. Melatonin concentrations.** Time course of melatonin concentrations during the first (solid line) and second (dashed line) evening in the laboratory in each condition (A "background", B "blue-dim", C "yellow-bright"). We show the mean with error bars representing the standard error.

**Visual Comfort (S4).** There was moderate (inconclusive) evidence against a difference between the conditions regarding visual comfort experienced during the light exposure (i.e., average rating on 5-item Likert scales on how pleasant/bright/glaring the light was, and how pleasant the light colour was). The data was approximately 6 times more likely under the H0 than under the H1 ($BF_{10}$ = 0.16). For the condition × time interaction, there was moderate evidence in favour of H1 ($BF_{10}$ = 7.68). Supplemental Figure 1 shows the time course of the visual comfort ratings.

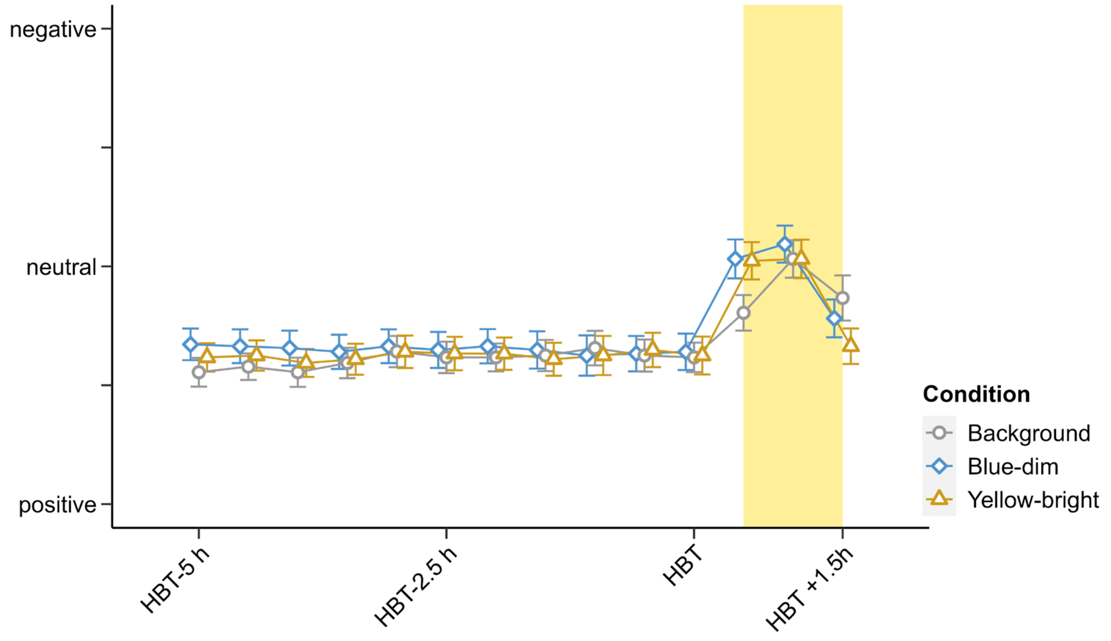


**Supplemental Figure 2: Visual comfort ratings.** Mean visual comfort scores during the first evening in the laboratory. Error bars indicate the standard error. The yellow box indicates the period of the light exposure. HBT = Habitual bedtime.

**PVT: Median Reaction Time (RT; S5), Fastest 10% RTs (S6), Slowest 10% RTs (S7).** Analyses revealed extreme evidence in favour of a condition difference during the light exposure with the data being approx. 153 times more likely under the H1 than the H0 ($BF_{10}$ = 153.33). More precisely, there was conclusive evidence in favour of faster median reaction times in the background ($mean_{background}$ = 384.4 ± 35.5 ms) than the yellow-bright ($mean_{yellow\text{-}bright}$ = 395.5 ± 32.6 ms; $BF_{10}$ = 29.06) or the blue-dim condition ($mean_{blue\text{-}dim}$ = 399.8 ± 39.1 ms; $BF_{10}$ = 260.1). There was only anecdotal (inconclusive) evidence against a difference between the yellow-bright and the blue-dim conditions ($BF_{10}$ = 0.39). There was anecdotal evidence against a condition × time interaction with the data being approx. 2 times more likely under the H0 compared to H1 ($BF_{10}$ = 0.47).

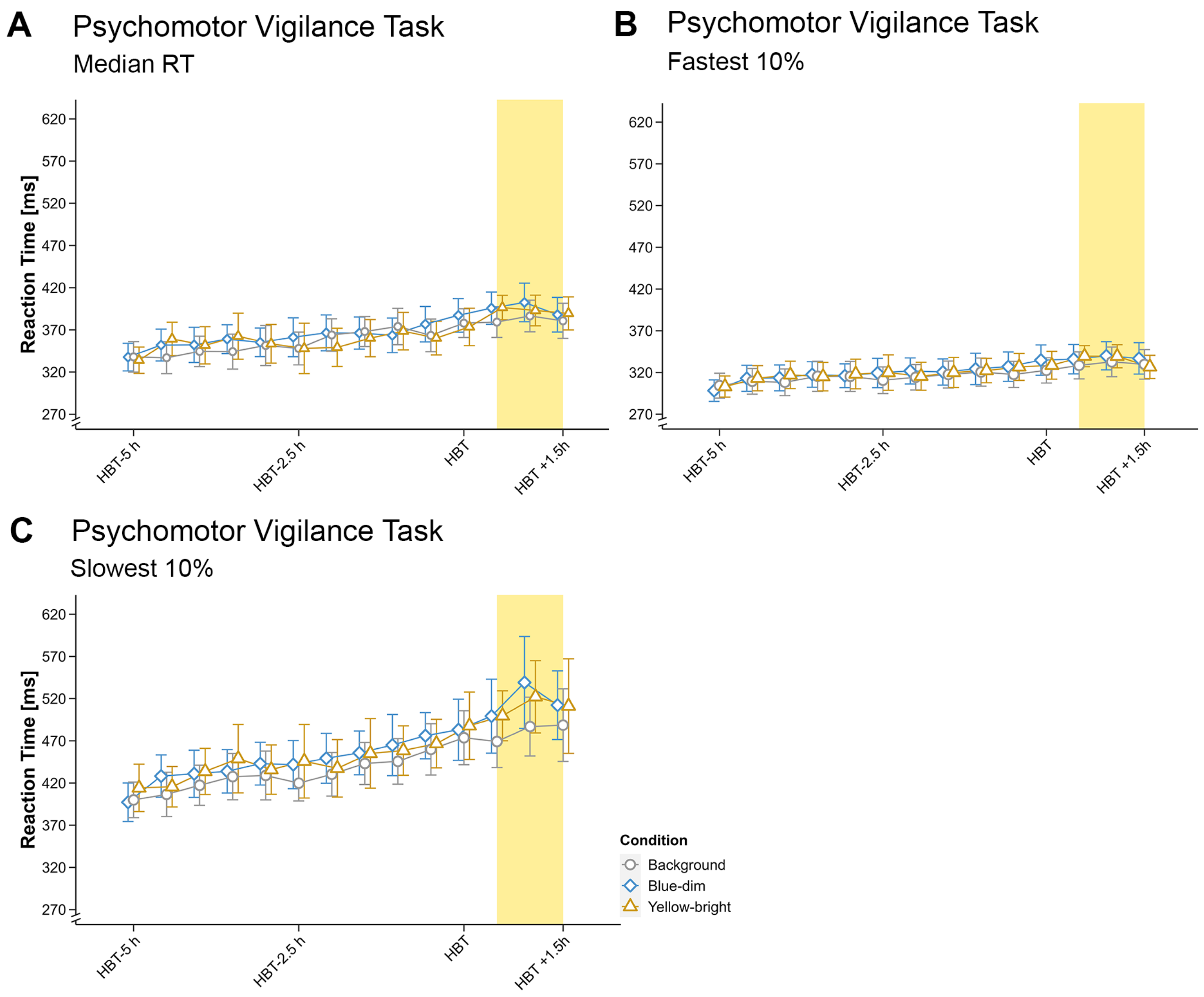


**Supplemental Figure 3: Reaction times on the psychomotor vigilance task (PVT).** A Median, B mean of the 10% fastest, C mean of the 10% slowest reaction times across the first evening in the laboratory. Error bars indicate 95% confidence intervals. The yellow box indicates the duration of the light exposure. HBT = Habitual bedtime.

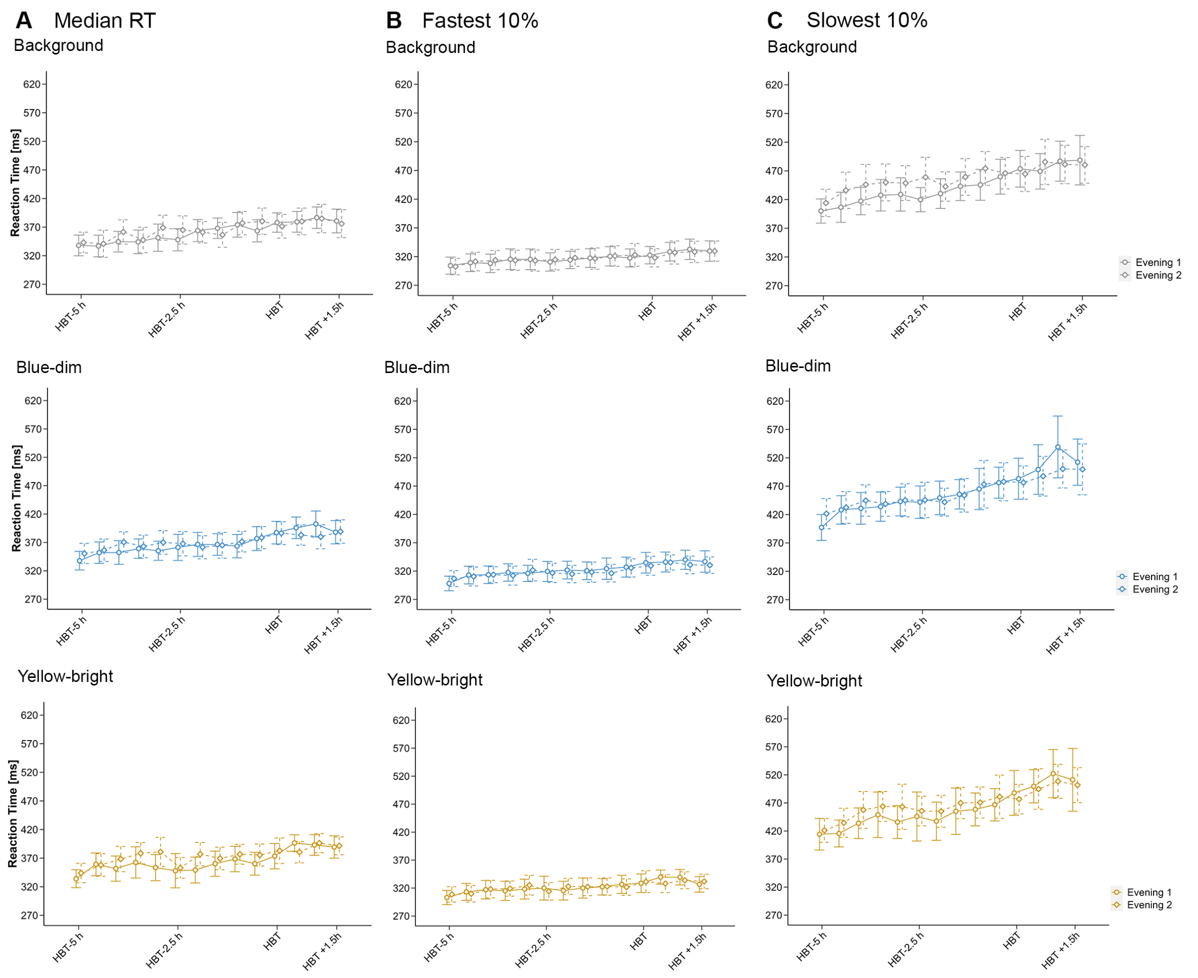


**Supplemental Figure 4: Reaction times on the psychomotor vigilance task (PVT).** A Median, B mean of the 10% fastest, C mean of the 10% slowest reaction times across the first (solid lines, circles) and second (dashed lines, diamonds) evenings in the laboratory. Error bars indicate 95% confidence intervals. Note that the data from the second evening were not part of the analysis plan and are only shown here for completeness. We thus also refrain from statistical analyses. HBT = Habitual bedtime.

**EEG-derived Sleep Onset Latency (SLAT; S8).** Analyses yielded inconclusive evidence against a condition difference regarding the onset latency to 10 minutes of continuous sleep. The data were approx. 3 times more likely given the H0 than the H1 ($BF_{10} = 0.31$; $BF_{10\ \text{rank-based}} = 0.39$). The mean latency to 10 minutes of continuous sleep was 17.0 ± 42.2 minutes in the background, 9.4 ± 13.2 minutes in the yellow-bright, and 7.5 ± 4.3 minutes in the blue-dim condition.

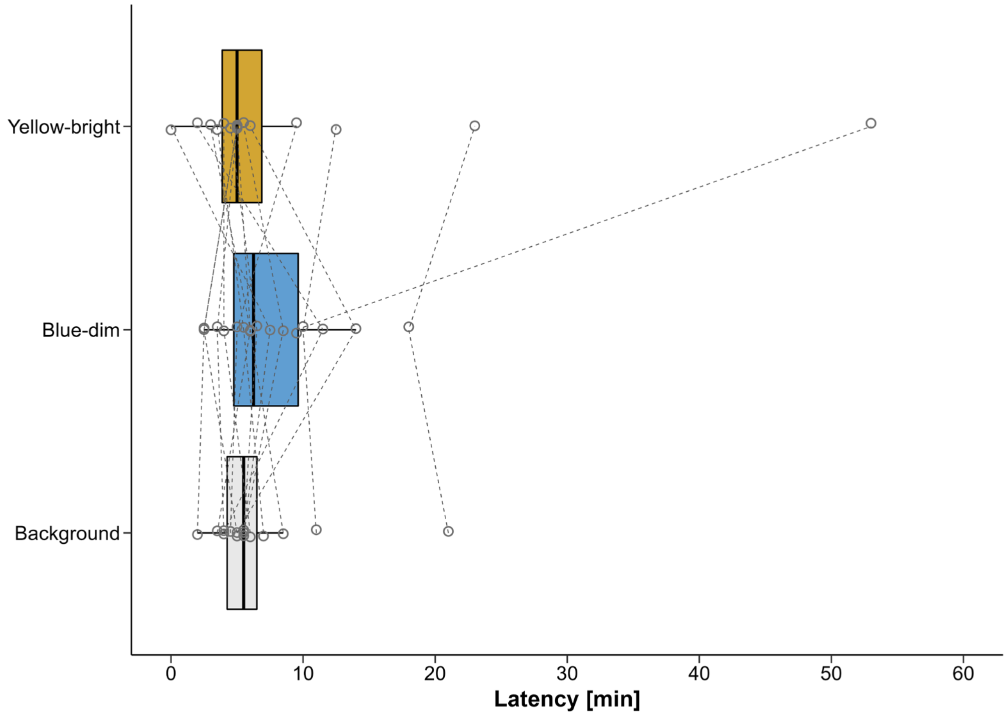


**Supplemental Figure 5: Latency to 10 minutes of continuous sleep.** In the boxplots, the lower and upper hinges of the boxplot correspond to the 25% and 75% quartiles, the thick black line indicates the median. Whiskers extend to the lowest/largest value at most 1.5× the interquartile range (IQR) from the hinges. Gray circles represent individual values of participants. Note: The individual data point from one participant, who had a latency of 174.5 min to 10 min of continuous sleep in the background condition, was removed from the plot as this would have concealed the pattern of the other data points.

#### Subjective Sleepiness (S2).

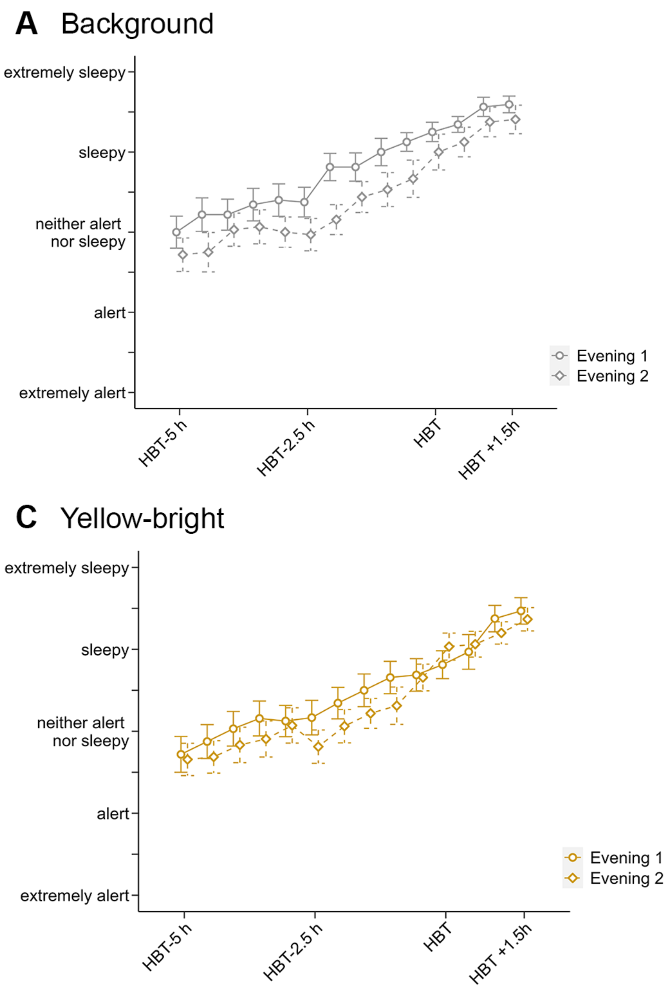


**Supplemental Figure 6: Subjective Sleepiness.** Mean and standard error of subjective sleepiness as assessed with the Karolinska Sleepiness Scale (KSS) during evening 1 (solid lines, circles) and evening 2 (dashed lines, diamonds) in the three light exposure conditions. Note that the data from the second evening were not part of the analysis plan and are only shown here for completeness. We thus refrain from statistical analyses. HBT = Habitual bedtime.

#### Supplemental Methods

##### Protocol.

The reason to not fully randomise the order of conditions was that a full randomisation would have required us to acquire data from multiples of 12 participants (i.e., always 6 men and 6 women). Given our financial and resource limit of 16 participants, this would not have worked out.

**Statistical data analysis.**

For all analyses, we report $BF_{10}$, i.e., the likelihood of the data under H1 compared to H0. We tested for normality using Shapiro-Wilk's test and for homogeneity of variances across the three conditions using Levene's test. Levene's test was never significant, i.e., homogeneity of variances was assumed in all cases. Using QQ-Plots and plots of standardized residuals versus fitted values, we assessed the model fit using raw and rank-transformed values. In cases, where rank-transformation improved the fit, we report results for both analyses. Following general recommendations[1], error percentages below 20% were deemed acceptable (default: 10 000 iterations) as this should result in the same qualitative conclusion. Following general recommendations[1], error percentages below 20% were deemed acceptable (default: 10 000 iterations) as this should result in the same qualitative conclusion.

**Data acquisition.** Data acquisition took place between March and December 2022 with a break in August. More precisely, the following number of volunteers (in brackets) had their first experimental visit in each month: March (1), April (2), May (0), June (2), July (2), September (1), October (3), November (3), December (2). The Laboratory Log provides even more detailed information on when each participant had his/her visits: https://doi.org/10.6084/m9.figshare.23578695.

**Subjectively reported light history.** The following table gives an overview (mean ± SD; range) of the reported light history on the day the volunteers came to the lab.

**Supplemental Table 1.**

| | **Time under open sky [in hours]** | **Light quality** | **Amount of light** |
|---|---|---|---|
| **Visit 1** | 1.6 ± 1.2 (0.5-4.5) | 6.4 ± 2.7 (1-10) | 6.9 ± 2.3 (2-10) |
| **Visit 2** | 1.5 ± 1.3 (0.3-5.3) | 6.8 ± 2.3 (1-10) | 5.2 ± 1.8 (2-8) |
| **Visit 3** | 1.7 ± 1.5 (0.5-5.3) | 6.1 ± 2.4 (3-10) | 5.8 ± 2.1 (3-10) |

*Comments*. Light quality and the amount of light that reached the eyes was assessed using Likert scales (range 0-10; 0 = very dull day/ very little light, 10 = bright summer day/ a lot of light).

#### Supplemental Tables

##### Suppl. Table 2. Sleep Descriptives.

| | Sleep Onset Latency [min] | Latency to 10 min continuous sleep [min] | Sleep Efficiency [%] | Wake after Sleep Onset [min] | Number of Awakenings | N1 Latency [min] | N1 Percent | N2 Latency [min] | N2 Percent | N3 Latency [min] | N3 Percent | REM Latency [min] | REM Percent |
|---|---|---|---|---|---|---|---|---|---|---|---|---|---|
| **Background** | 5.8±2.4 | 19.6±46.8 | 92.6±10.5 | 20.7±36.7 | 10.9±5.0 | 5.8±2.4 | 6.9±2.9 | 8.7±2.7 | 44.6±4.8 | 34.0±44.0 | 24.8±6.6 | 111.2±73.6 | 23.7±4.6 |
| **Blue-dim** | 5.7±3.0 | 7.5±4.2 | 92.9±6.4 | 19.5±21.4 | 9.9±5.5 | 5.7±3.0 | 6.3±3.3 | 8.5±3.8 | 44.8±6.0 | 23.3±7.3 | 24.4±6.0 | 96.7±49.6 | 24.5±6.9 |
| **Yellow-bright** | 3.9±1.9 | 10.0±14.2 | 94.8±3.9 | 14.7±12.9 | 11.2±7.0 | 3.9±1.9 | 6.8±2.9 | 7.4±4.1 | 45.0±5.2 | 26.1±17.6 | 24.9±6.2 | 104.2±78.3 | 23.3±7.2 |
| **Visit 1** | 5.8±2.4 | 19.6±46.8 | 92.6±10.5 | 20.7±36.7 | 10.9±5.0 | 5.8±2.4 | 6.9±2.9 | 8.7±2.7 | 44.6±4.8 | 34.0±44.0 | 24.8±6.6 | 111.2±73.6 | 23.7±4.6 |
| **Visit 2** | 5.7±2.7 | 8.0±5.2 | 93.2±6.1 | 18.5±19.9 | 11.5±6.4 | 5.7±2.7 | 7.1±3.1 | 8.8±4.1 | 44.0±4.9 | 23.5±8.0 | 24.5±5.5 | 85.5±44.4 | 24.5±5.6 |
| **Visit 3** | 3.9±2.4 | 9.5±13.9 | 94.5±4.5 | 15.7±15.3 | 9.6±6.0 | 3.9±2.4 | 6.0±2.9 | 7.0±3.7 | 45.8±6.1 | 25.8±17.4 | 24.8±6.6 | 115.5±78.5 | 23.4±8.3 |

Participants had a 6-h sleep opportunity starting 2 hours after habitual bedtime. Values reflect the mean ± the standard deviation. Note that Visit 1 was always the "Background" condition.

### References

1 van Doorn, J. *et al.* The JASP guidelines for conducting and reporting a Bayesian analysis. *Psychonomic Bulletin & Review* **28**, 813-826 (2021). https://doi.org/10.3758/s13423-020-01798-5
